# Ecological specialization promotes diversity and diversification in the East Mediterranean genus *Ricotia* (Brassicaceae)

**DOI:** 10.1101/2020.08.30.274670

**Authors:** Barış Özüdoğru, Çağaşan Karacaoğlu, Galip Akaydın, Sadık Erik, Klaus Mummenhoff, İsmail Kudret Sağlam

## Abstract

Despite its amazing biodiversity, the Eastern Mediterranean remains a highly understudied region especially when compared to the Western Mediterranean. Scarcity of such studies restrict our understanding of the processes shaping diversity across the entire Mediterranean. To this end we used a combination of molecular markers and presence/absence data from all species of the Eastern Mediterrranean genus *Ricotia* collected across its full geographic range to determine historical, ecological and evolutionary factors responsible for lineage-specific diversification in this genus. Network analysis based on nuclear ribosomal and chloroplast DNA revealed high genetic structure within all lineages and phylogenetic reconstructions based on the multispecies coalescent showed that within lineage diversification corresponded to the onset of the Mediterranean climate. Reconstruction of ancestral histories indicate the genus originated within Anatolia and slowly spread across the Eastern Mediterranean and Levant using the Taurus mountains. Ecological niche models based on climatic and environmental variables suggest local populations did not go through any major distributional shifts and have persisted in present day habitats since the LGM. Furthermore, niche differentiation tests revealed significant niche differences between closely related species and showed the main variables predicting species limits to be different for each species. Our results give crucial information on the patterns and processes shaping diversity in the Eastern Mediterranean and show the main factor promoting diversification to be local environmental dynamics and ecological specialization and not large scale latitudinal movements as often reported for southern Europe. By determining regional and global patterns of diversification in an eastern Mediterranean genus we further our understanding of the major trends influencing plant diversity in the Mediterranean basin as a whole.

## 1. Introduction

The Mediterranean Basin is one of the most diverse floristic regions in the world, hosting around 25,000 plant species, 60% of which are endemics (Nieto Feliner, 2014; Euro+Med PlantBase, 2016; Escudero et al., 2018). Although there are several studies aimed at understanding the processes and patterns contributing to this outstanding diversity, a large majority of these studies focus on the Western Mediterranean region (Escudero et al., 2018 and references therein). In contrast very little attention has been paid to eastern Mediterranean groups more specifically those around Anatolia and adjacent regions (Özüdoğru et al., 2015). The scarcity of these studies restrict our understanding of the dynamics and processes that have shaped patterns of divergence across the Mediterranean region as a whole.

Located at the crossroads between the Mediterranean, Irano-Anatolian and Caucasus hotspots and with about 10,000 plant species, Anatolia is one of the most floristically diverse regions in the World (Davis, 1965; Güner et al., 2012; Gür, 2016). Hosting over 700 crucifer taxa (Güner et al., 2012), Anatolia is also widely regarded as the center of origin for the family Brassicaceae (Franzke et al., 2009). Additionally, as a part of the Alpine-Himalayan orogenic belt Anatolia has a high level of topographic and climatic diversity with a wide range of habitats stretching from sea level to 5000 meters (Şekercioğlu et al., 2011). These unique features make Anatolia a prime candidate for understanding the dynamics of the Eastern Mediterranean region and furthering our understanding of the Mediterranean basin as a whole.

It is well known that, with the exception of high mountain peaks, Anatolia was free of glaciation during the Pleistocene and lowlands were covered by forest and steppe communities (Ansell et al., 2011; Şenkul and Doğan, 2013). Therefore Anatolia is widely regarded as an important refugium for temperate species during the Pleistocene (Atalay, 1996; Çiplak et al., 2010). Despite Anatolia’s potential importance, there are relatively few studies exploring the regions impact on diversification of plant taxa at the species or genus level (e.g. Ansell et al., 2011; Koch et al., 2016; Smýkal et al., 2017; Özüdoğru et al., 2020). What studies there are, concentrate mostly on the role of Anatolia as a refugium and center of diversity for widespread taxa that later recolonized Europe (Ansell et al., 2011; Ali et al., 2016; 2019) and not on local forces that drive and shape diversification within Anatolia or the Eastern Medditerenean at large. In conclusion, Anatolia can be regarded as a biogeographically exciting but under-explored region (Ansell et al., 2011) with no study to date providing a comprehensive breakdown of historical, evolutionary and ecological factors shaping diversification of a East Mediterranean genus and all its species.

*Ricotia* is a small crucifer genus with 10 species, restricted to the Eastern Mediterranean region (Fig. 1). Though small in species number, the genus provides an excellent opportunity to map the phylogeographic and evolutionary dynamics of Eastern Mediterranean lineages as a whole since it includes annual vs. perennial, insular vs. mainland, lowland vs. high alpine, and Irano-Turanian vs. Mediterranean species (Özüdoğru et al., 2015; Özçandir et al., 2019).

**Fig. 1.**
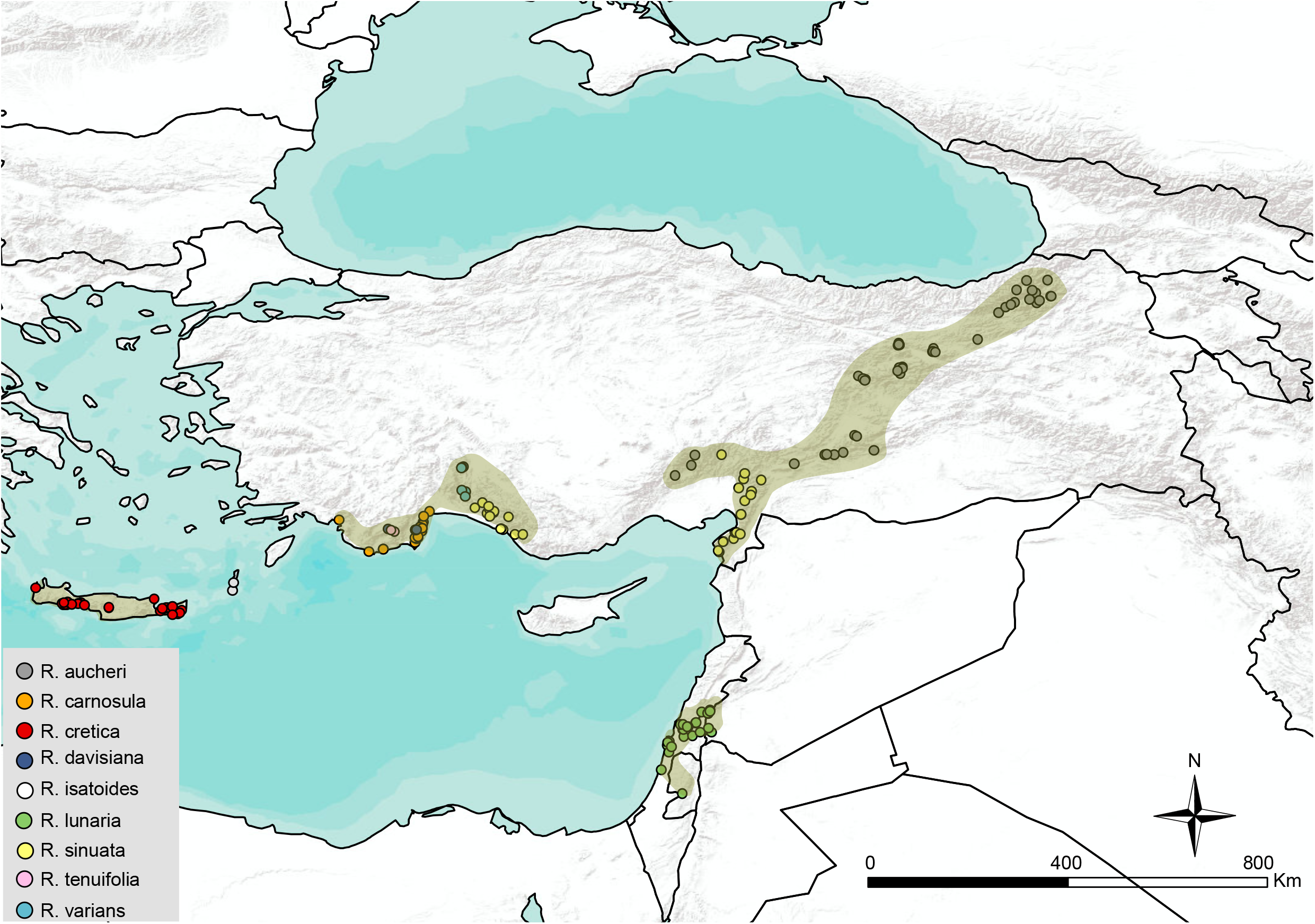
Distribution of *Ricotia* species/accessions used in the current study.

The biogeographical history of *Ricotia* was already described by Özüdoğru et al. (2015) based on ITS and *trn*L-F markers but this study did not investigate the influence of Eastern Mediterranean dynamics on the complex demographic and evolutionary history of *Ricotia* species and its ecotypes or tackle questions regarding the origin of the genus. Using additional genetic markers and accessions, the current study aims to clarify the origin of the genus and together with ecological niche modeling determine historical, ecological and evolutionary factors responsible for lineage-specific diversification patterns of *Ricotia* species. We show that regional and global patterns of diversification in the genus *Ricotia* is driven by local environmental dynamics and ecological specialization and not by large scale events like contraction-expansion in and out of glacial refugia. By determining regional and global patterns of diversification in an eastern Mediterranean genus we further our understanding of the major trends influencing plant diversity and diversification in the Mediterranean basin as a whole.

## 2. Material and methods

### 2.1. Plant sampling

To cover the full extent of morphological variation and complete distributional area of *Ricotia* species we collected samples from 65 distinct populations, representing nine *Ricotia* species. Samples were either collected directly from the field or from relevant herbariums and seed banks (Supplementary material, Table S1). In total we sequenced 156 individuals (1-4 per population) at two different loci (ITS and *trnL-F* + *trn*Q-5’ rps16; hereafter cpDNA) using standard methods (for details see supplementary material). To this data set we added 4 sequences of the tribe Biscutelleae (*Biscutella laevigata, Heldreichia bupleurifolia, Lunaria rediviva* and *Megadenia pygmaea*) to serve as an outgroup (Özüdoğru et al., 2015; 2017). Voucher specimens of representative individuals have been deposited at Hacettepe University Herbarium (HUB) and new sequences unique to this study have been uploaded to the GenBank database (Accession No: 7068956). A recently described tenth *Ricotia* species, *R. candiriana*, was left out of the main study because we could only obtain two ITS sequences from this species. Phylogenetic placement of *R. candiriana* based on ITS sequences is given in Supplementary material, Fig. S1.

### 2.2. Genetic diversity and haplotype analyses

For each data set, genetic variation was quantified by calculating haplotype (Hd) and nucleotide (Θ□ and ΘS) diversity. Departures from neutrality were assessed using Tajimas’s D (Tajima, 1989), and Fu’s Fs (Fu, 1997) tests. All estimations were done in DNASP, version 5.10 (Librado and Rozas, 2009) and confidence intervals were determined using coalescent simulations.

CpDNA haplotype and ITS ribotype networks were constructed by PopArt (Leigh and Bryant, 2015). Prior to analyses, repeat regions in the cpDNA data set (e.g. TATTATAT, TAAAATA in *trn*L-F, or TTGTATATAGT in *trn*Q-5-*rps*16) were eliminated from the haplotype description process. Alignment gaps were considered as a 5th character state in the haplotype definition process but some of the accessions (e.g. one population of *R. sinuata* (H49), from Yaylaalani district, Antalya region Turkey) were eliminated or masked in the cpDNA data set due to including an ~200 bp deletion in the *trn*Q-5’ rps16 region. Networks were constructed separately for each of the four *Ricotia* species (*R. aucheri, R. carnosula, R. cretica* and *R. sinuata*), the three main *Ricotia* lineages (Aucheri clade, Tenuifolia clade and core *Ricotia* clade) and for the genus as a whole. Six local *Ricotia* species with restricted distribution and low individual sampling were left out of the analysis.

### 2.3. Historical demography

Population expansion in the four *Ricotia* species (*R. aucheri, R. carnosula, R. cretica* and *R. sinuata*) was tested using pairwise differences between individual haplotypes/ribotypes under the sudden expansion model of Rogers and Harpending (1992) as implemented in DnaSP version 5.10 (Librado and Rozas, 2009). Significance of the sudden expansion model was assessed by calculating the raggedness index (r) (Harpending, 1994) and Ramos-Onsins and Rozas’ test statistic (R2) (Ramos-Onsins and Rozas, 2002).

Effective population size change through time was further evaluated using extended Bayesian skyline plots (EBSP) (Heled and Drummond, 2008). EBSP can analyze multiple unlinked loci simultaneously and therefore, improve upon demographic inferences from single-locus analyses. EBSP analysis was conducted using a strict clock model and calibrated using the per lineage mutation rate of each gene estimated from the dated phylogeny reconstructed in the *BEAST analysis. Two independent runs with 50 million generations, and sampling every 5000 generations were conducted to check for convergence and the first 10% of the genealogies were discarded as burn-in.

### 2.4. Phylogenetic analyses and time estimations

To understand evolutionary relationships and divergence times between *Ricotia* species we estimated a time calibrated species tree based on both ITS and cpDNA data sets using the multispecies coalescent procedure *BEAST (Heled and Drummond 2010). Prior to analyses, the best fitting model of substitution for each marker was selected using the corrected Akaike Information Criterion (AICc) as performed in MEGA6 (Tamura et al., 2013). Divergence times were estimated based on an uncorrelated lognormal relaxed clock model and trees were calibrated by setting the crown age of tribe Biscutelleae and genus *Ricotia* on the cpDNA tree to 21.65 ±1 and 17.8 ±1 mya respectively as given in Mandakova et al. (2018) under an uncorrelated lognormal relaxed clock model. A calibrated Yule process of speciation was used as a tree prior for the species tree, while a constant population size coalescent model was used for inferring the relationships within species. Phylogenetic and dating analyses were performed in BEAST v.2.6.1 (Bouckaert et al. 2014).

We conducted two independent MCMC runs with chain lengths of 50 million, sampling every 5000 generations. 10% of each run was discarded as burn-in and convergence between independent runs (ESS values > 500) was checked using tracer, version 1.5 (http://tree.bio.ed.ac.uk/software/figtree) and combined using logcombiner version 1.7.1. Consensus (maximum clade credibility, MCC) gene and species trees with divergence times were summarized from this posterior distribution using treeannotator, version 1.7.1, and visualized using figtree, version 1.3.1 (http://tree.bio.ed.ac.uk/software/figtree) and DensiTree 2 (Bouckaert and Heled, 2014).

### 2.5. Historical biogeography

To determine the biogeographic history of the genus we reconstructed ancestral ranges of *Ricotia* species under six different models based on dispersal–extinction–cladogenesis (DEC, DEC+J), dispersal–vicariance (DIVALIkE, DIVALIKE+J) and Bayesian inference of historical biogeography (BayArea, BayArea+J), with +J adding founder□event speciation to each model (Matzke, 2013, 2014).

Ancestral state reconstruction under all models was based on a seperate cpDNA tree estimated using only unique haplotypes to minimize the influence of uneven sampling numbers within groups (i.e. species). A time calibrated cpDNA tree with only unique haplotypes was estimated in BEAST under the same parameter values given above with the only difference being that we used a constant population size coalescent model as a tree prior instead of a Yule process.

Biogeographic regions were defined following Özüdoğru et al. (2015) as: (A) islands of the southern arc of Tertiary folded mountains including Crete, Karpathos and Rhodes, (B) Western part of south Anatolia (= Antalya section), (C) Eastern part of south Anatolia (= Adana section), (D) Anatolian Diagonal Mountains, and (E) Israel/Lebanon. Models were compared using the corrected Akaike information criterion (AICc) and the best fitting model/models were used to describe the biogeographic history of the genus. All analyses were conducted using the R package BioGeoBEARS (Matzke, 2013; Matzke, 2014) as implemented in RASP (Yu et al., 2015).

### 2.6. Ecological niche modelling (ENM)

To reconstruct the present and past distribution of five relatively widespread *Ricotia* species; *R.aucheri, R.carnasula, R.cretica, R.sinuata* and *R.lunaria*, we generated ecological niche models using the presence-only maximum entropy algorithm as implemented in MaxEnt 3.4.1 (Phillips et al., 2006; Elith et al., 2011). In order to reduce issues related to background selection we limited the area of interest for ENMs to the known distribution of each species (Elith et al., 2011; Merow et al., 2013, Brown et al., 2016).

ENMs were based on a total of 160 occurrence records pertaining to the 5 species obtained from field studies and present-day bioclimatic, spatial heterogeneity and soil classification GIS data layers at 30 arc-seconds (~1 km) resolution. 19 bioclimatic variables along with altitude, slope and aspect data were obtained from the WorldClim database (Hijmans et al., 2005; www.worldclim.org). Spatial heterogeneity was quantified using 14 metrics based on the Enhanced Vegetation Index (EVI) acquired by the Moderate Resolution Imaging Spectroradiometer (MODIS) (Tuanmu and Jetz, 2015; https://www.earthenv.org/texture) (Supplementary material, Table S2-3). Soil classification data containing The World Reference Base (WRB)/FAO classes (layers coded as TAXNWRB) was obtained from the ISRIC World Soil Information SoilGrids portal (http://soilgrids.org).

Using MAXENT we built two present day distribution models, one using all environmental variables (dataset 1) and the second using only the 19 bioclimatic variables (dataset 2). Prior to analysis we performed a paired Pearson correlation test to remove highly correlated parameters allowing for a maximum correlation coefficient of 0.8 (Feldman et al., 2017). To predict distribution of species at the Last Glacial Maximum (LGM) we projected models obtained from our present day species-bioclimatic analysis (dataset 2) onto the LGM using three general circulation models (CCSM4, MIROC-ESM, and MPI-ESM-P) available at WorldClim. Paired Pearson correlation test was performed to identify and remove highly correlated parameters used in each model. Correlation test performed using SDMtoolbox version 1.1c (Brown, 2014) and allowing maximum 0.8 correlation coefficient (Feldman et al., 2017). Accuracy of model predictions were evaluated using the area under the (receiver operating characteristic) curve (AUC) calculated in MAXENT (Yi et al., 2016; Fois et al., 2018).

Niche differentiation between the 5 *Ricotia* species was determined in ENMTools 1.3 using niche overlap and identity tests (Warren et al. 2010). Pairwise comparisons for niche overlap and identity between all species was based on Schoener’s D and Hellinger’s I (Warren et al., 2008) which ranges from 0 (complete divergence/no overlap) to 1 (high similarity/complete overlap).

## 3. Results

### 3.1. Diversity and historical demography

The aligned ITS and cpDNA data matrices included 56 ribotypes and 70 haplotypes (plus four representatives of tribe Biscutelleae) respectively (for detailed statistics see Supplementary material, Tables S4-S5). For the four *Ricotia* species, ribotype and haplotype diversity was highest in *R. aucheri* and lowest in *R. carnosula* (in ITS) and *R.cretica* (in cpDNA) and in all four species Tajima’s D and Fu’s Fs values were negative, but none of them were statistically significant (Supplementary material, Tables S6-S7). Haplotype and ribotype networks of the four *Ricotia* species showed a distinct east-west disjunction in both *R. cretica* and *R. sinuata* whereas no such pattern was evident in *R. aucheri* or *R. carnosula* even though the former showed high structuring in both ribo and haplotype diversity (Fig. 2, supplementary material Figs. S2-S5). Our network analysis also indicated that the Fethiye population of *R. carnosula* was well separated from the main groups by 30 mutational steps.

**Fig. 2.**
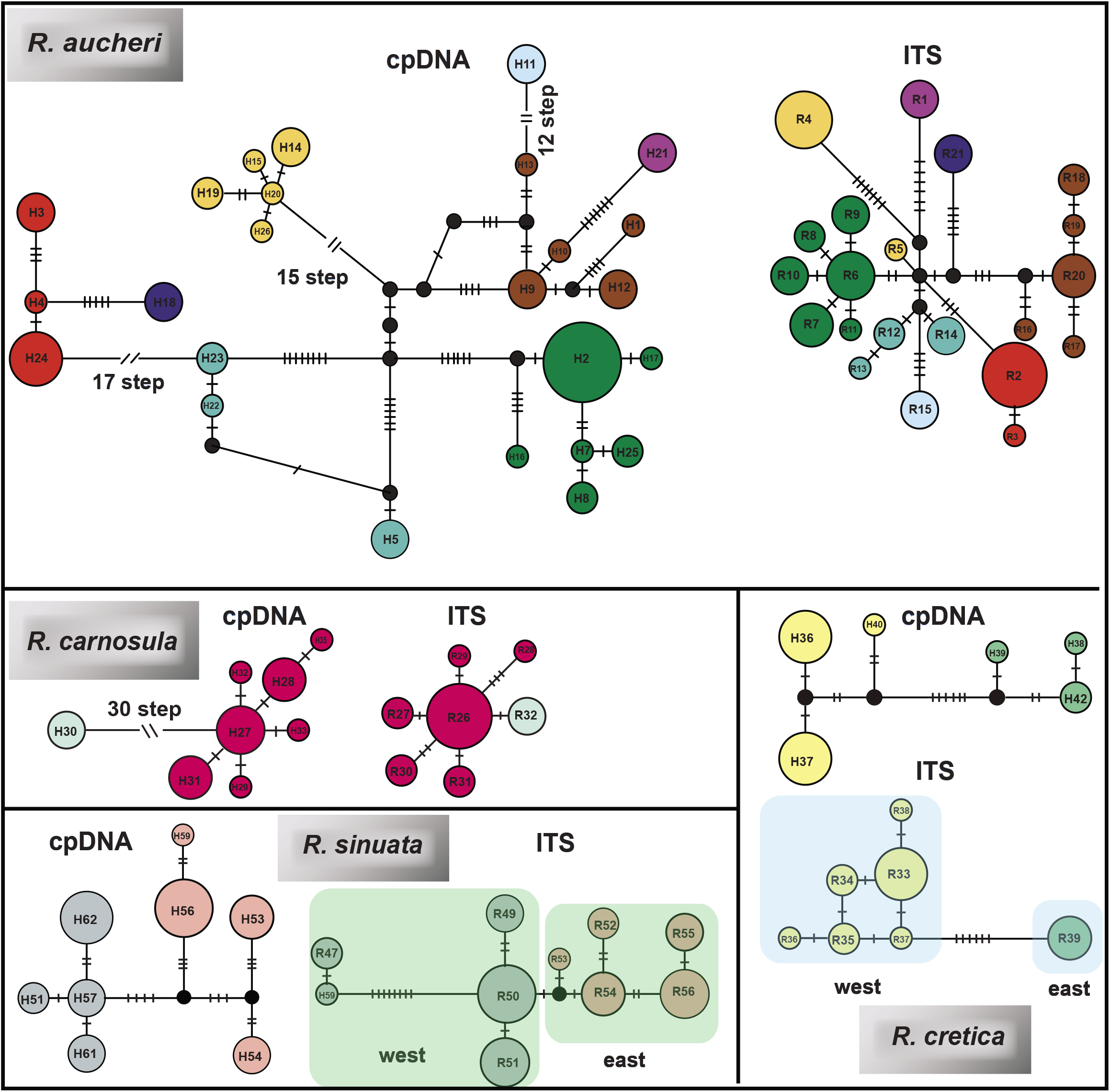
Haplotype (cpDNA) and ribotype (ITS) networks of four *Ricotia* species used in demographic and Ecological Niche Modelling analyses (*R. aucheri, R. carnosula, R. sinuata, R. cretica*)

Mismatch distributions and EBSP analyses showed no sign of recent or historical population expansion and indicated that all species have generally remained stable since 0.5 Ma, although the *R. aucheri* showed a slight decline towards the present (Fig. 3).

**Fig. 3.**
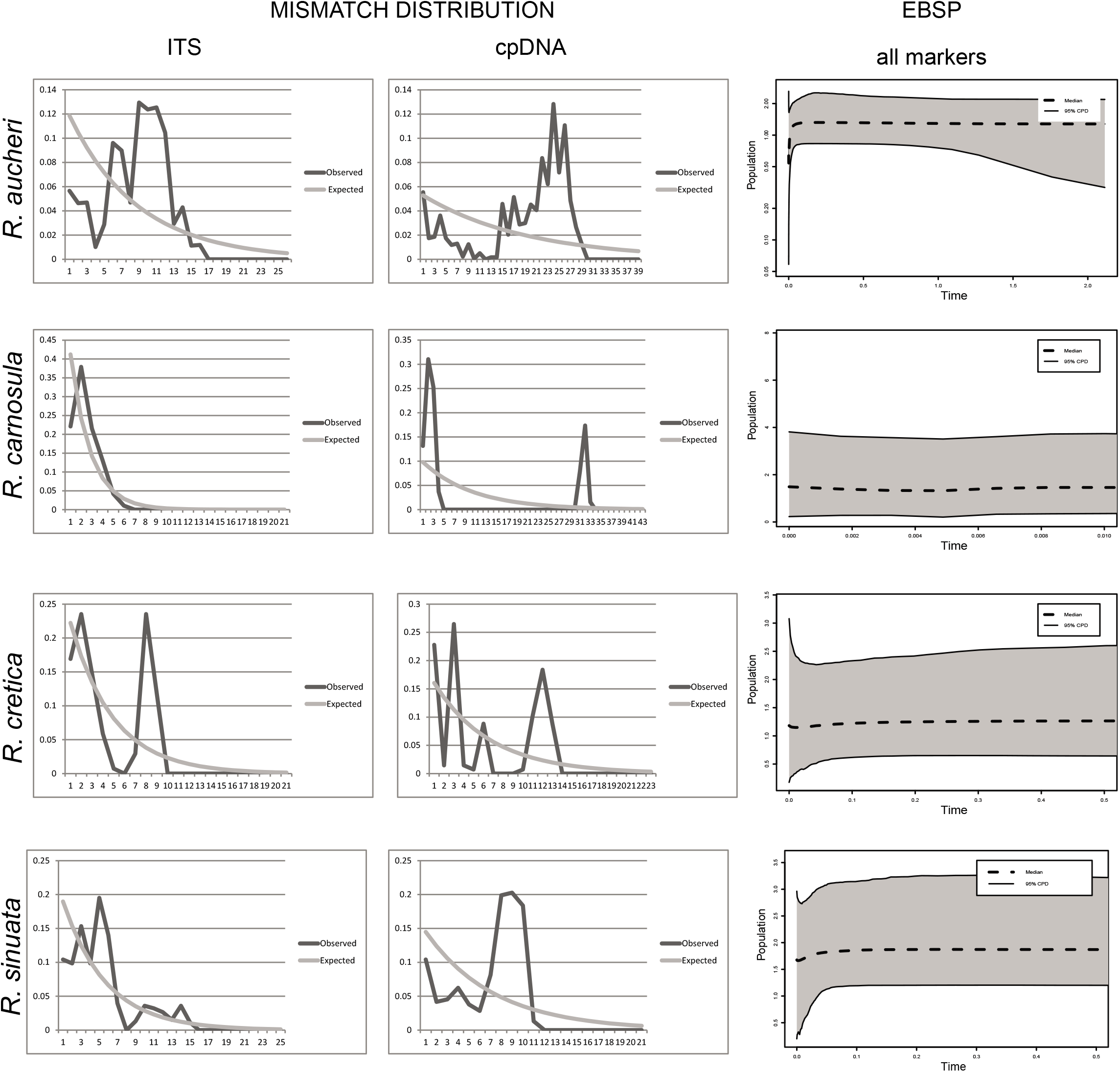
Demographic analysis of *R. aucheri, R. carnosula, R. cretica*, and *R. sinuata*. Graphical outputs of mismatch distribution analyses of pairwise haplotype differences based on ITS and cpDNA data and Expanded Bayesian Skyline Plots (EBSP) based on all markers.

### 3.2. Phylogeny and time estimation

Multispecies coalescent analysis based on ITS and cpDNA regions resulted in a well resolved species tree able to separate all three *Ricotia* lineages with high accuracy (Fig. 4). The species tree placed the Aucheri group as the basal clade with Tenuifolia and the main *Ricotia* lineage forming sister clades. Divergence time estimates indicate that seperation of all three lineages was complete around 16 mya (14.7-19.1) and that within lineage diversification (i.e. speciation) occured after the onset of the Mediterranean Climate (3.4-2.8 Mya).

**Fig. 4.**
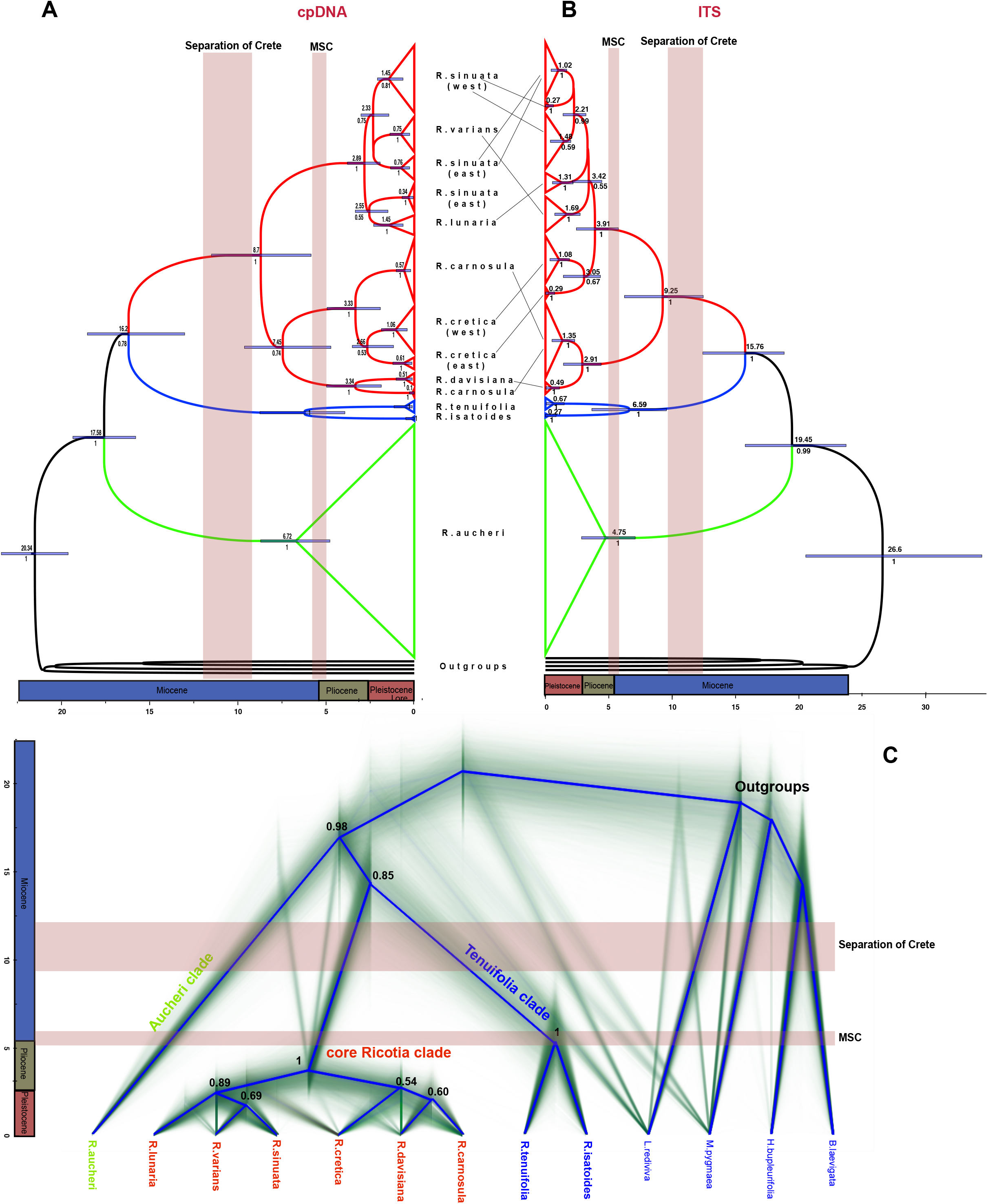
Gene trees and species tree of *Ricotia* based on *Beast analysis of all markers (ITS + cpDNA). Representatives of tribe Biscutelleae (*Biscutella laevigata, Heldreichia bupleurifolia, Lunaria rediviva* and *Megadenia pygmaea*) served as outgroup. Shown is the complete genealogical tree set (thin lines) and consensus tree (thick lines in blue) visualized by DensiTree. Posterior probability values are given at nodes only for *Ricotia* species (ingroup).

Although the species tree was well resolved at the lineage level, support values for species within the main *Ricotia* lineage were low indicating high uncertainty for relationships between these taxa. This uncertainty can be traced back to differences in the position of species or populations groups within the made *Ricotia* clade for the cpDNA and ITS gene trees. The ITS gene tree supported monophyly of both *R. sinuata* and *R. carnosula* whereas both species were found to be paraphyletic in the cpDNA gene tree. In addition, placement of *R. cretica* lineages differed between the two gene trees. The ITS gene tree placed both *R.cretica* lineages together with the *R.lunaria, R.sinuata* and *R.varians* clades whereas in the cpDNA gene tree *R.cretica* can be found together with the *R. carnusula* and *R. davisiana* clades. Finally, as suggested in Özçandir et al. (2019), the recently described *R. candiriana* fell into the same clade with its sister species *R. davisiana* (Supplementary material, Fig. S1)

### 3.3. Historical biogeography

Based on the model test implemented in BioGeoBEARS, the DEC +J model resulted in the lowest corrected AIC value (85.53) and Ln L= 39.59 (Supplementary material, Table S8). The DEC +J model suggests the Antalya region (region B) as the most probable ancestral area of the genus *Ricotia* as a whole (Fig. 5, P = 0.75) and indicates that most lineages diverged within this region, including the Mediterranean *Ricotia* species (core clade + *Tenuifolia* clade, P=98.28), the *Tenuifolia* Clade (P=90.13) and the clade including both the Crete endemic *R. cretica* and the mainland species *R.davisiana* and *R. carnosula* (P=99.44). Our analysis also showed that *R. lunaria* distributed around Israel originated from eastern southern Anatolia (P=75.33) (for details see Supplementary material, Table S9)

**Fig. 5.**
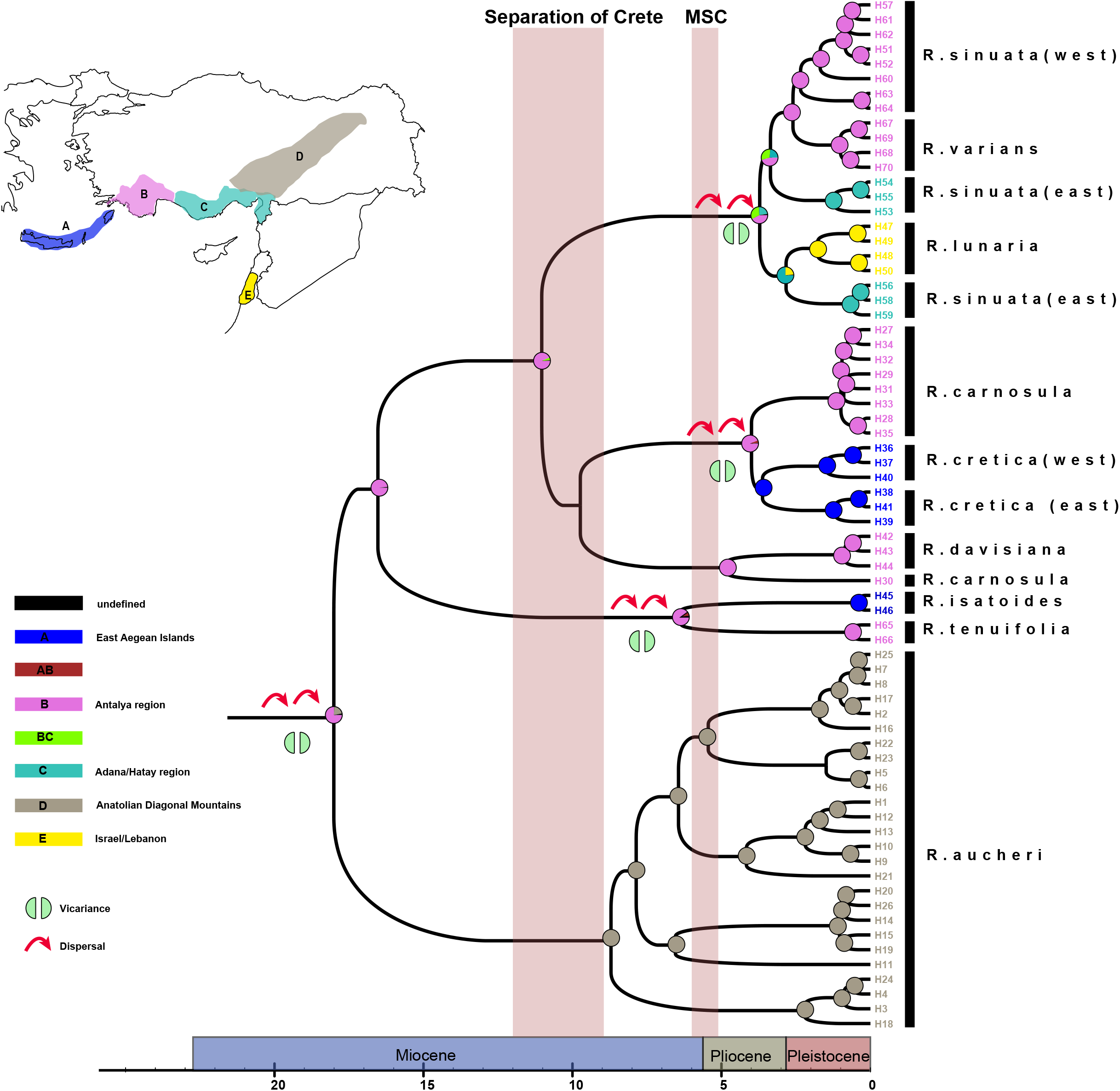
Ancestral area reconstruction for *Ricotia* using the DEC+j model in BioGeoBEARS based on the BEAST-derived maximum clade credibility (MCC) tree of the cpDNA data set (Figure S1 in Supporting Information). The insert map shows the geographical distribution of *Ricotia* species according to Özüdoğğu et al (2015). The colour code defines extant and possible ancestral ranges and specific symbols refer to vicariance (two green semi-circles) and dispersal (red arrow) events, respectively. The relative probability of each alternative ancestral range for nodes is shown in pie charts.

### 3.4. Ecological niche modelling (ENM)

Models for current distribution of species obtained from data set 1 and 2 showed high predictive power with good to very good average AUC values. For *R. aucheri* and *R. carnosula* predictions based on data set 1 (all environmental variables) were a much better fit to the known distribution of these species (Fig. 6). For these species non-climatic variables like normalized dispersion of the Enhanced Vegetation Index (cv-EVI) and soil (taxnwrb) were found to be the most influential factors shaping distribution (Supplementary material, Table S10). For *R. cretica, R.lunaria and R.sinuata*, data set 2 (only climatic variables) had higher predictive power (Supplementary material, Table S10) and the climatic variables slope, precipitation seasonality (BIO15) and annual precipitation (BIO12) were the most influential factors shaping their distribution (Table S10). Projections of the current ENMs to the three GCMs suggested little or no change between the current distribution of the species and that at the LGM except for *R. carnosula* which seems to have been restricted to the Levant region during the LGM (Figure 6).

**Fig. 6.**
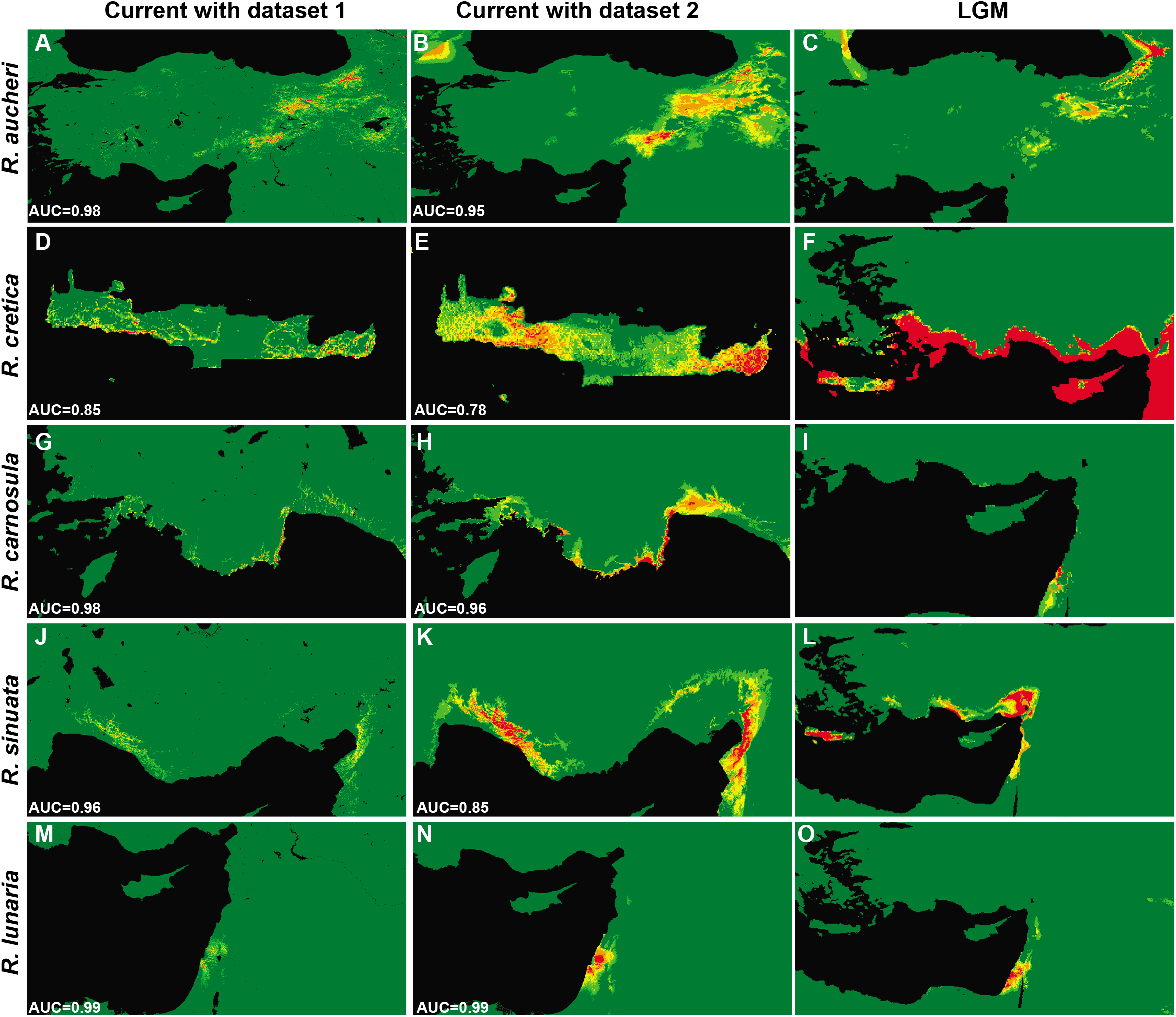
Results of species distribution modelling for *R. aucheri, R. carnosula, R. cretica, R. sinuata*, and *R. lunaria* using the maximum entropy algorithm in the eastern Mediterranean (sub)regions as implemented in Maxent. Analyses are based on environmental variables, and projected to past conditions at the last glacial maximum (LGM, c. 21 kya). The first column is based on the results of all variables (dataset 1) whereas the second column represents the results of only climatic variables (dataset 2). Marginal to optimal habitats are shown in green (marginal) to red (optimal).

Niche differentiation tests, performed through ENMTools returned significant differences between environmental niches of all species indicating that species utilized distinct environmental spaces. Observed pairwise niche overlap (D) values under the current environmental conditions (data set 2) was below 0.2 (Table 1) for all species supporting little to no overlap. Pairwise identity tests showed significant differences between all comparisons and rejected the null hypothesis of niche identity in all cases (Table 1).

## 4. Discussion

Relationships between the three main *Ricotia* lineages were well resolved on the species tree and in agreement with previously published results (Özüdoğru et al., 2015; Mandakova et al., 2018). Divergence time estimates indicate that the three main *Ricotia* lineages split fairly close to the MRCA of the genus (~17.8 mya) and diverged from another well before major geological events like the separation of Crete and the Messinian Salinity Crisis (Kougioumoutzis et al., 2018). Although phylogenetic relationships between the three main *Ricotia* lineages were well resolved on the species tree, there was appreciable uncertainty between species within the core *Ricotia* lineage due to discrepancies between ITS and cpDNA gene trees. Discrepancies between gene trees in terms of the position of species or population groups of the core *Ricotia* clade might be the consequence of deep historical range shifts. As an example, the unexpected position of the Fethiye population of *R. carnosula* and the close relationships between *R. lunaria* and eastern clade of *R. sinuata* could be explained by historical range shifts pre-dating the LGM and not captured by our ENM models.

Our reconstruction of ancestral history provides some evidence for such past migrations as it highlights a series of migratory events along with chromosomal rearrangements (as already shown by Mandakova et al., 2018) resulting in the spread and diversification of the three main lineages. Based on our biogeographical analyses, the most recent common ancestor of *Ricotia* originated in the Antalya region and from there spread northeast to the Irano-Anatolian area establishing *R. aucheri* by 6 mya, and later towards southeast and southwest establishing *R. cretica* in Crete by 4 mya and *R. lunaria* in the Levant by 2 mya. Therefore, despite deep divergence of the three main lineages, within lineage diversification was comparably recent and corresponded to the Plio/Pleistocene and onset of the Mediterranean climate (3.4-2.8 mya). Radiation of ancestral *Ricotia* populations from the Mediterranean into the Irano-Anatolian region after the onset of the Mediterranean climate is also fully compatible with the paleogeographic history of the region as much of the Anatolian Diagonal and surrounding regions were covered by the Tethys Sea up until the late Miocene (~ 12 mya) (Erinç, 1953; Şengör, 1980; Bartol and Govers, 2014).

The current distribution of *Ricotia* species in Anatolia matches the recently described Taurus Way (Çiplak, 2008; Kaya and Çiplak, 2017). This corridor connects North-East Anatolia to Western Anatolia and the Balkans via the Anatolian Diagonal and Taurus Mountains and therefore is an important dispersal route for mountainous species. In addition to being a dispersal corridor, high altitude species utilizing the Taurus way can also shift their distribution to lower altitudes under suitable conditions which may present opportunities for in-situ diversification and ecological specialization. Life history patterns observed for *Ricotia* species along the Taurus way agree well with this assumption. It has already been established that perenniality in *Ricotia* species is ancestral and several shifts to an annual life cycle has occurred in the genus (Özüdoğru et al., 2015). Accordingly, ancestral perennial forms of *Ricotia* species occupy high mountainous habitats along the Taurus and East Anatolia mountains whereas their annual relatives inhabit habitats at lower elevations along the southern Anatolian coast.

Therefore, similar to many Mediterranean plant lineages (Tremetsberger et al., 2016; Benítez-Benítez et al., 2018; Alonso et al., 2020) climatic oscillations in the Pleistocene and the onset of the Mediterranean climate seems to have driven both radiation and diversification in these Eastern Mediterranean lineages. However, contrary to northern and central European lineages the dynamics southern taxa do not correspond to the classical contraction and post-glacial expansion model but seems to be the result of in-situ divergence caused by ecological specialization. In the absence of large glaciations as witnessed in Northern Europe, taxa indigenous to the southern Mediterranean region could survive climatic oscillations of the Quaternary via short distance altitudinal shifts to more favorable habitats (Sağlam et. al. 2014). As a result, while the classical contraction-expansion paradigm (Hewitt, 1996) can serve as a robust model for explaining observed patterns in widespread woody or herbaceous plant species (Taberlet et al., 1998; Nieto Feliner, 2014) it can not be generalized to southern European lineages which are influenced more by complex local dynamics (Nieto Feliner, 2014; Ali et al., 2016, 2019).

There are several lines of evidence in the present study that support this assertion. Firstly, in all lineages both haplotype and ribotype networks indicate complex intraspecific histories (Fig. 2) and none of them show a star-shaped network, which are classical indicators of recent expansion out of refugia (Avise, 2000). Secondly, ENM results indicate that local populations did not go through any major distributional shifts and have for the most part persisted in present day habitats since the LGM (Figs. 6 and S6). This scenario is also supported by multimodal mismatch distributions and stable effective population sizes of populations in Bayesian skyline plots which indicate little to no demographic change throughout the Pleistocene (Fig. 3). Lastly, our ENM models revealed significant ecological niche differentiation between species and showed that the main variable(s) for predicting (or explaining) distributional limits of species was different for each species. In conclusion, diversification within this group seems to be driven by local dynamics and ecological specialization and to the best of our knowledge, this is the first quantitative study reporting ecological vicariance for a plant genus throughout the Aegean Islands, Anatolia and the Levant.

Furthermore, ecological specialization can also explain both the east-west divide observed in *R. cretica* and *R. sinuata* and the highly structured distribution of *R. aucheri* despite the presence of any substantial geographic barriers. Our ENM analysis supports this conclusion as the current distribution of all species were shaped mostly by environmental variables like slope and precipitation seasonality for *R. cretica*, annual precipitation for *R.sinuata* and vegetation cover for *R. aucheri*.

For *R. cretica*, a distribution gap in the central part of Crete can be seen in both present and past predictions (Fig. 5D-F) indicating that this distinction between east and west has been in place since the LGM. East and western populations of *R. cretica* also show important phenological differences. While individuals in West Crete were still flowering and just started fruiting in May, eastern *R. cretica* plants were totally mature and already dead at this time (personal observation, BÖ). East-West disjunctions on Crete were documented for various other organisms most notably in trap-door spiders where again climatic differentiation between east and western Crete was proposed to be responsible for the observed divide (Thanou et al., 2017). Thus, we tentatively conclude that local adaptation could have played an important role in genetic isolation between eastern and western populations of *R. cretica*.

A similar East-West disjunction was observed in *R. sinuata* in the cpDNA gene tree (Fig. 4), although all accessions in the ITS analyses were monophyletic. ENM analysis for *R. sinuata* and *R. lunaria* indicates that the two species might have been sympatrically distributed during the LGM (Fig. 6). Therefore, east west patterns in *R. sinuata* might also be explained by haplotype sharing (i.e. hybridization) between *R. sinuata* and *R. lunaria* and might not necessarily indicate ecological specialization.

Like other scree plants *R. aucheri* is highly restricted in its distribution due to the fragmented nature of its habitats. Throughout its distribution along the Taurus and Anatolian Diagonal Mountains, *R. aucheri* co-exist with other typical scree plants like *Heldreichia bupleurifolia* (monotypic genus *Heldreichia*, Brassicaceae) in high alpine plant communities (Parolly et al., 2010; Özüdoğru et al., 2015). But contrary to *H. bupleurifolia* which does not show any biogeographic structuring *R. aucheri* is highly structured and can be split into distinct haplo-/ribotype clusters representing different local populations along the Anatolian Diagonal (Supplementary material, Fig. S2). The main environmental gradient that causes the Anatolian Diagonal to serve as a steep ecological barrier for the distribution of species has been recognized as “temperature seasonality” (Gür, 2016). Likewise, temperature seasonality as well as “vegetation type/cover” and “annual precipitation” were the main factors explaining the present day distribution of *R. aucheri* (Table S10). This again suggests that genetic structuring of this species along the Anatolian diagonal results from ecological isolation rather than a physical barrier and emphasizes that the Anatolian Diagonal does not need to serve only as a physical barrier between east and west.

## CONCLUSION

The Mediterranean region has long been the focal point of many phylogeographic studies mainly in regards to its role as a refugium for the preservation of genetic diversity throughout glacial advances (Médail & Diadema, 2009; Coello et al., 2020). However, much less attention has been given to the complex phylogeographic structures observed in this region and the myriad of processes that can lead to diversity across such varied topographies. One possible reason for this bias is that most studies from the Mediterranean region have concentrated on well known southern refugia like Iberia and Italy which have undoubtedly played a very important role in the re-colonization of central and northern Europe (Nieto Feliner, 2014). Here we present crucial information on the patterns and processes shaping diversity in the Eastern Mediterranean and show that as a result of the reduced effects of climatic oscillations in the region, diversification of eastern lineages have largely occurred insitu over a long span of time instead of large latitudinal movements. Therefore, the main factor promoting diversification in this region seems to be ecological specialization as structured populations adapt to different conditions spread across the patchy landscapes of the Mediterranean.

Recent studies from the Western Mediterenean have also noted similarly complex phylogeographic patterns and an overall lack of general models explaining divergence across the Mediterenean basin (Lo Presti and Oberprieler, 2011; Fernández-Mazuecos and Vargas, 2011; Benítez-Benítez et al., 2018). In conclusion the contraction-expansion dynamics often cited to explain diversity patterns within central and northern Europe is not a good representative for understanding the dynamics of local Mediterranean flora where divergence and radiation seem to be driven by long term ecological adaptation and specialization. Therefore, a clear distinction should be made between the dynamics of non-indegenous northern flora that use the southern Mediterranean as a refuge during glacial periods and those of local flora which have for the most part remained stable throughout the Quaternary.

## Supporting information

Suppementary Information

## Acknowledgements

We thank Golshan Zare, Hasan Yildirim and M.Yavuz Paksoy for providing *Ricotia* samples, and curators of B, E, G and HUB for making their herbarium material available for this study. This study was supported by Hacettepe University Scientific Research Projects Coordination Unit (Projects no. 011D03601002, 01301704001 and FBI-2018-17147) and is the part of a Ph.D. thesis by the first author at Hacettepe University.

## Author contributions

BÖ and KM conceived study. BÖ, GA, SE and KM collected samples. BÖ performed the molecular work and analyzed data with input from IKS. ÇK performed ENM analyses. BÖ and IKS wrote the manuscript with contributions from ÇK, and KM.

## Notes

### Competing Interest Statement

The authors have declared no competing interest.

### Summary of Updates

Supplementary files have been revised

