## Supplementary material for "Ecological specialization promotes diversity and diversification in the East Mediterranean genus *Ricotia* (Brassicaceae)": Suppementary Information: Figure S1.pdf

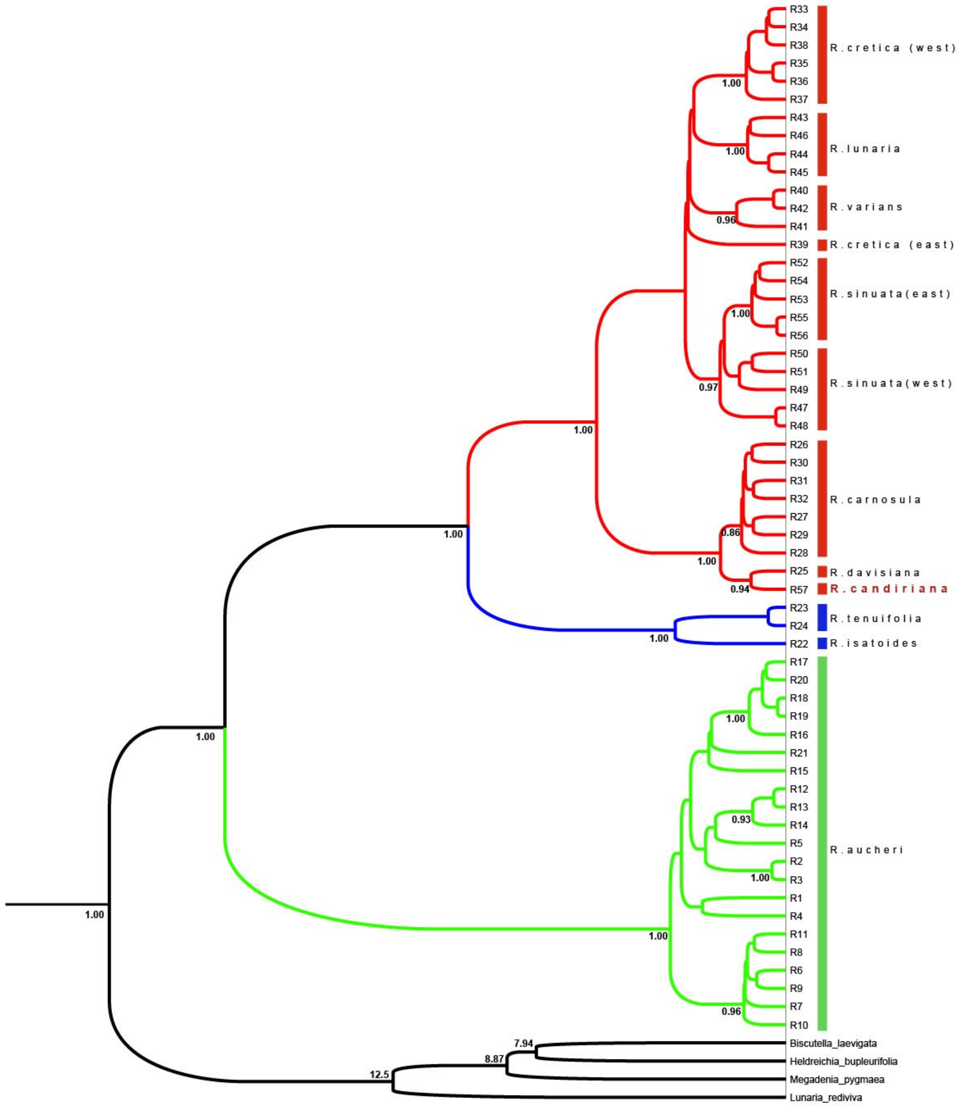


**Figure S1** Bayesian phylogenetic tree of *Ricotia* ribotypes including recently described *Ricotia candiriana*. Bayesian consensus tree with posterior probability values (PP) > 0.5. Bayesian PP values and mean values of node ages are given at nodes. Blue node bars indicated the 95% confidence interval for of the node age estimate. Green, blue and red colors indicate the three *Ricotia* clades (Aucheri clade, Tenuifolia clade and main *Ricotia* clade respectively) based on the studies of Özüdoğru et al.,( 2015) and Mandakova et al. (2018).
