## Supplementary material for "Ecological specialization promotes diversity and diversification in the East Mediterranean genus *Ricotia* (Brassicaceae)": Suppementary Information: Supplementary methods.docx

**Supplementary methods S1**

**DNA EXTRACTION, POLYMERASE CHAIN REACTION, SEQUENCING, AND SEQUENCE ALIGNMENT**

Total genomic DNA was extracted from silica dried leaf material, herbarium samples or fresh materials from germinated seeds using DNeasy Plant Mini Kit (Qiagen, Hilden, Germany) following the manufacturer’s instructions.

ITS-1 and ITS-4 (White *et al.* 1990) primers were used for amplifying the nuclear ribosomal ITS region (ITS1‒5.8S‒ITS2), c and f (Taberlet et al., 1991) for *trnL*(UAA) intron/*trnL-trnF* intergenic spacer (herafter *trn*L-F), and forward and reverse for *trnQ-*5’*rps16* (Shaw et al., 2007). Purification and sequencing were performed by Biyoeksen (İstanbul, Turkey).

**ECOLOGİCAL NİCHE MODELLİNG (ENM)**

Different data sets were used to estimate the appropriate areas for species current distribution. First, 19 bioclimatic variables (detailed information about these parameters can be found at www.worldclim.org/bioclim) and altitude data are obtained from WorldClim database (Hijmans, Cameron, Parra, Jones, & Jarvis, 2005; www.worldclim.org). Slope and aspect data are derived from altitude data downloaded from WorldClim. Second dataset was Global Habitat Heterogeneity dataset which contains 14 metrics (Coefficient of variation, Evenness, Range, Shannon, Simpson, Standard deviation, Contrast, Correlation, Dissimilarity, Entropy, Homogeneity, Maximum, Uniformity, Variance) quantifying spatial heterogeneity of global habitat based on the textural features of Enhanced Vegetation Index (EVI) acquired by the Moderate Resolution Imaging Spectroradiometer (MODIS) (Tuanmu & Jetz, 2015; https://www.earthenv.org/texture). Another data was soil classification data which is downloaded from ISRIC World Soil Information SoilGrids portal (http://soilgrids.org). The downloaded soil classification data contains The World Reference Base (WRB)/FAO classes and named as TAXNWRB. All data mentioned above is downloaded at 30 arc-seconds (~1 km) resolution and used ENM analyses as “dataset 1”.

**Bioclimatic variables used in this study**

BIO1 = Annual Mean Temperature

BIO2 = Mean Diurnal Range (Mean of monthly (max temp - min temp))

BIO3 = Isothermality (BIO2/BIO7) (* 100)

BIO4 = Temperature Seasonality (standard deviation *100)

BIO5 = Max Temperature of Warmest Month

BIO6 = Min Temperature of Coldest Month

BIO7 = Temperature Annual Range (BIO5-BIO6)

BIO8 = Mean Temperature of Wettest Quarter

BIO9 = Mean Temperature of Driest Quarter

BIO10 = Mean Temperature of Warmest Quarter

BIO11 = Mean Temperature of Coldest Quarter

BIO12 = Annual Precipitation

BIO13 = Precipitation of Wettest Month

BIO14 = Precipitation of Driest Month

BIO15 = Precipitation Seasonality (Coefficient of Variation)

BIO16 = Precipitation of Wettest Quarter

BIO17 = Precipitation of Driest Quarter

BIO18 = Precipitation of Warmest Quarter

BIO19 = Precipitation of Coldest Quarter

The first order and second order texture measures derived from EVI composite (Tuanmu & Jetz, 2015). The first order texture measures are statistics describing the frequency distribution of pixel values in a pixel neighborhood within an image. Second-order measures are based on the probability of observing a pair of values at two pixels with a given inter-pixel distance and orientation (Tuceryan & Jain, 1998).

Hijmans, R. J., Cameron, S.E., Parra, J. L., Jones, P. G., & Jarvis, A. (2005). Very high resolution interpolated climate surfaces for global land areas. *International Journal of Climatology*, 25(15), 1965–1978. https://doi.org/10.1002/joc.1276

Shaw, J., Lickey, E. B., Schilling, E. E., & Small, R.L. (2007). Comparison of whole chloroplast genome sequences to choose noncoding regions for phylogenetic studies in angiosperms, the Tortoise and the Hare III. *American Journal of Botany*, 94, 275-288.

Taberlet, P., Gielly, L., Pautou, G. & Bouvet, J. **(**1991). Universal primers for amplification of three non-coding regions of chloroplast DNA. *Plant Molecular Biology,* 17, 1105–1109. <http://dx.doi.org/10.1007/BF00037152>

Tuceryan, M., Jain, A.K. (1998) Texture analysis. The handbook of pattern recognition and computer vision (ed. by C.H. Chen, L.F. Pau and P.S.P. Wang), pp. 207–248. World Scientific, Singapore.

Tuanmu, M. N., & W. Jetz. (2015). A global, remote sensing-based characterization of terrestrial habitat heterogeneity for biodiversity and ecosystem modeling. *Global Ecology and Biogeography*. DOI: 10.1111/geb.12365.

White, T.J, Bruns, T., Lee, S. & Taylor, J. (1990). Amplification and direct sequencing of fungal ribosomal RNA genes for phylogenetics. *In*: Innis, M.A., Gelfand, D.H., Sninsky, J.J. & White, T.J. (Eds.) *PCR protocols: A guide to methods and applications*. Academic Press, New York, pp. 315–322. <https://doi.org/10.1016/B978-0-12-372180-8.50042-1>
