## Supplementary material for "Ecological specialization promotes diversity and diversification in the East Mediterranean genus *Ricotia* (Brassicaceae)": Suppementary Information: Table S1.docx

**Table S1.** Collection information for samples used in this study.

|  | **Species** | **Voucher (herbarium code)** | **Locality, altitude, coordinate (if possible) and collection date.** | **Number of Samples** | **Haplotypes**  **and Ribotypes** |
| --- | --- | --- | --- | --- | --- |
| 1 | *R. aucheri* | B. Özüdoğru 3561 & E.Çilden (HUB) | Turkey, Adıyaman: Nemrut Mountain, 12.5 km from Cendere to Kıran Village, screes, 960-970 m. Latitude: 37.979083 N, Longitude: 38.688383 E, 13.06.2014. | 3 | H21 / R1 |
| 2 | *R. aucheri* | B. Özüdoğru, 3353 (HUB) | Turkey, Erzincan: Kemaliye, around Apçağa Village, screes, 1370-1400 m. Latitude: 39.237821 N, Longitude: 38.52252 E, 07.06.2012. | 4 | H3, H4 / R2, R3 |
| 3 | *R. aucheri* | B. Özüdoğru 3352 (HUB) | Turkey, Erzincan: Kemaliye, 800-900 m from Sandık Village junction, screes, 965-1000 m, Latitude: 39.285468 N, Longitude: 38.479813 E, 07.06.2012. | 3 | H24 / R2 |
| 4. | *R. aucheri* | B. Özüdoğru 3350 (HUB) | Turkey, Erzincan: Kemaliye, above Sandık Bağı Village, Sulutarla, screes, 1100 -1150 m. Latitude: 39.271379 N, Longitude: 38.491389 E, 07.06.2012. | 3 | H24 / R2 |
| 5 | *R. aucheri* | MYP Tunceli-t (HUB) | Turkey, Tunceli Ovacık, on road Karagöl Plateau, 1800 m, Latitude: 39.415 N, Longitude: 39.117 E | 3 | H19, H20 / R4 |
| 6 | *R. aucheri* | Ş. Yıldırımlı 2298 (HUB) | Turkey, Tunceli: Ovacık, Bellihasan crest, Munzur Mountains, 2500-2700 m, 29.07.1979. | 1 | H26 / R5 |
| 7 | *R. aucheri* | H. Yıldırım 3101 (HUB) | Turkey, Tunceli:Ovacık, Aksu dere, around Kırkmar divan, screes, 2200 m, Latitude: 39.455056 N, Longitude: 39.19575 E, 20.07.2014. | 4 | H14, H15 / R4 |
| 8 | *R. aucheri* | A. Güner 4760 (HUB) | Turkey, Artvin: between Artvin and Yusufeli arası, Binatınbaşı, road sides, stony places, 350 m, 20.05.1983. | 1 | H2 / R6 |
| 9 | *R. aucheri* | A. Tatlı 4255 (HUB) | Turkey, Erzurum: Çoruh Valley, around Çamlıkaya, on andesite rocks, 1050 m, 14.05.1976. | 1 | H2 / R7 |
| 10 | *R. aucheri* | B. Özüdoğru 3569 (HUB) | Turkey, Erzurum: between Yusufeli and Tortum arası, from Morkaya Village to Tortum Lake, 2 km before Lake, screes, 815-830 m, Latitude: 40.68375 N, Longitude: 41.6707 E, 18.06.2014. | 2 | H25 / R8 |
| 11 | *R. aucheri* | B. Özüdoğru 3570 (HUB) | Turkey, Artvin: between Yusufeli and Olur, around Ayvalı Village, screes, 810-820 m, Latitude: 40.752167 N, Longitude: 41.888267 E, 18.06.2014. | 3 | H7, H8 / R9 |
| 12 | *R. aucheri* | B. Özüdoğru 3562 (HUB) | Turkey, Erzurum: 13 km from İspir to Yusufeli, roadsides, screes, 1035 m, Latitude: 40.556467 N, Longitude: 41.06225 E, 17.06.2014. | 3 | H2, H17 / R6 |
| 13 | *R. aucheri* | B. Özüdoğru 3563 (HUB) | Turkey, Erzurum: 24 km km from İspir to Yusufeli, roadsides, screes, 940 m, Latitude: 40.603107 N, Longitude: 41.160043 E, 17.06.2014. | 3 | H2 / R7 |
| 14 | *R. aucheri* | B. Özüdoğru 3565 (HUB) | Turkey, Erzurum: on the road of Barhal (Yusufeli), 2 km from Yusufeli to Dereiçi Village, roadside, eroded places, 710 m, Latitude: 40.861013 N, Longitude: 41.539853 E, 18.06.2014. | 3 | H2 / R6, R10 |
| 15 | *R. aucheri* | H. Yıldırım 3120 (HUB) | Turkey, Artvin: Yusufeli, Sarıgöl, on the road Yüksekoba Plateau, screes, 1580 m, Latitude: 41.040578 N, Longitude: 41.436417 E, 23.07.2014. | 1 | H16 / R10 |
| 16 | *R. aucheri* | G. Zare (HUB) | Turkey, Arvtin Yusufeli, Dikmenli Village, roadside, 512 m, Latitude: 41.046153, Longitude: 41.820109. | 1 | H2 / R11 |
| 17 | *R. aucheri* | B. Özüdoğru 3471 & G. Akaydın (HUB) | Turkey, Erzurum: Aşkale-Bayburt road, 16. km, screes, 1800 m, Latitude: 39.982222 N, Longitude: 40.561667 E, 13.07.2013. | 3 | H5, H6 / R12, R13 |
| 18 | *R. aucheri* | B.Özüdoğru 3472 & G.Akaydın (HUB) | Turkey, Erzincan: Sakaltutan Pass, screes, 2040 m, Latitude: 39.87986 N, Longitude: 39.15303 E, 13.07.2013. | 3 | H22, H23 / R14 |
| 19 | *R. aucheri* | B. Özüdoğru 3217 & K. Özgişi (HUB) | Turkey, Kahramanmaraş: between Çağlayancerit and Ağabeyli, 25 km to Ağabeyli, screes, 1420 m, Latitude: 37.736888 N, Longitude 37.249948 E, 30.07.2011. | 3 | H11 / R15 |
| 20 | *R. aucheri* | A. Güner 3638 (HUB) | Turkey, Adıyaman: between Yazıbaşı (Azıkhan) Village and Körtigerri, steppe, metamorphic and calk places, Latitude: 37.923697 N, Longitude: 38.161142 1200-1800 m, 27.05.1981. | 1 | H1 / R16 |
| 21 | *R. aucheri* | H. Yıldırım 2723 (HUB) | Turkey, Malatya: Doğanşehir, Erkenek, Akdağ, below summit, screes, 1826-1900 m, Latitude: 37.902917 N, Longitude 37.970722 E, 21. 06. 2013. | 1 | H13 / R17 |
| 22 | *R. aucheri* | B. Özüdoğru 3354 (HUB) | Turkey, Malatya: Beydağı, west side, screes, 2060-2200 m, Latitude: 38.2325 N, Longitude 38.381667 E, 13.06.2012. | 4 | H9, H10 / R18,R19,R20 |
| 23 | *R. aucheri* | B. Özüdoğru 3559 E. Çilden (HUB) | Turkey, Malatya: 11 km from Erkenek to Gölbaşı, eroded sides, screes, 1040 m, Latitude: 37.910652 N, Longitude 37.840011 E, 13.06.2014. | 3 | H12 / R20 |
| 24 | *R. aucheri* | B. Özüdoğru 3545 & E. Çilden (HUB) | Turkey, Kayseri: Yahyalı, Kapuzbaşı, Aladağ forest road, 14. km, screes, 1200 m, Latitude: 37.714404 N, Longitude 35.401102 E, 12.06.2014. | 3 | H18 / R21 |
| 25 | *R. isatoides* | B. Özüdoğru 3475 (HUB) | Greece, Karpathos Island, Aperi Village, screes, 450-470 m, Latitude: 35.555278 N, Longitude 27.164167 E, 26.07.2013. | 2 | H45, H46 / R22 |
| 26 | *R. tenuifolia* | B. Özüdoğru 2968 (HUB) | Turkey, Antalya: between Finike and Elmalı, 1 km to Avlanbeli Pass, roadside, rocks, 1075 m, Latitude: 36.539497 N, Longitude 29.989191 E, 20.05.2011. | 3 | H65 / R23, R24 |
| 27 | *R. tenuifolia* | Paksoy 1080 (HUB) | Turkey, Antalya: Antalya: between Finike and Elmalı, calcareous rocks, 390 m, 24.04.2010. | 1 | H66 / R23 |
| 28 | *R. davisiana* | B. Özüdoğru 3215 & K.Özgişi (HUB) | Turkey, Antalya: Kemer, Tahtalı Mountain, north side, around Çukuryayla, screes, 1400-1500 m, Latitude: 36.548056 N, Longitude 30.421944 E, 21.07.2011. | 3 | H43, H44 / R25 |
| 29 | *R. davisiana* | H. Peşmen 4619 & A. Güner (HUB) | Turkey Antalya: Kemer, Tahtalı Mountain, between Yaylakuzdere and Peynirlik Kızılalan, calcareous rocks, 800-1600 m, Latitude: 36.549066 N, Longitude 30.446771 E, 04.05.1979. | 1 | H42 / R25 |
| 30 | *R. carnosula* | B. Özüdoğru 2852 (HUB) | Turkey, Antalya: Beldibi, opposite of Grand Ring Hotel, under *Pinus brutia*, 18 m, Latitude: 36.690556 N, Longitude 30.571111 E, 28.04.2011. | 1 | H28 / R26 |
| 31 | *R. carnosula* | B. Özüdoğru 2862 (HUB) | Turkey, Antalya: 10 km from Tekirova to Kumluca, rocky slopes, under *P.brutia*, Latitude: 36.493889 N, Longitude 30.484167 E, 130-150 m, 28.04.2011. | 2 | H34, H35 / R26 |
| 32 | *R. carnosula* | B. Özüdoğru 2884 (HUB) | Turkey, Antalya: Kumluca, North-East of Adrasan village, calcareous rocks, 210–230 m, Latitude: 36.320849 N, Longitude 30.415408 E, 30.04.2011. | 2 | H27 / R27, R28 |
| 33 | *R. carnosula* | B. Özüdoğru 2883 (HUB) | Turkey, Antalya: around Olympos, macchie, 128 m Latitude: 36.393056 N, Longitude 30.443333 E, 30.04.2011. | 1 | H32 / R27 |
| 34 | *R. carnosula* | B. Özüdoğru 2863 (HUB) | Turkey, Antalya: 3 km from Çıralı to Chiemera, under *P.brutia*, slopes, 20 m, Latitude: 36.414722 N, Longitude 30.474722 E, 28.04.2011. | 2 | H27, H29 / R26, R29 |
| 35 | *R. carnosula* | B. Özüdoğru 3437 & G. Akaydın (HUB) | Turkey, Antalya: Kemer, on the road of Tahtalı cable car station, road side, under *P.brutia*, 100 m, Latitude: 36.533017 N, Longitude 30.542894 E, 19.05.2013. | 2 | H27, H33 / R26 |
| 36 | *R. carnosula* | B. Özüdoğru 3315 (HUB) | Turkey, Antalya: Kas, around peninsula, macchie, 22 m, Latitude: 36.199907 N, Longitude: 29.631875 E, 14.04.2012. | 4 | H31 / R30, R31 |
| 37 | *R. carnosula* | B. Özüdoğru 3316 (HUB) | Turkey, Muğla Fethiye, above Yanıklar Village, under *P. brutia*, 70–80 m, Latitude: 36.738252 N, Longitude: 29. 064588 E, 25.04.2012.ş | 3 | H30 / R32 |
| 38 | *R. carnosula* | B. Özüdoğru, 2857 (HUB) | Turkey, Antalya: around Kemer, under *P. brutia*, on serpentine rocks, 37 m, Latitude: 36.581389 N, Longitude 30.546111 E, 28.04.2011. | 3 | H28 / R26 |
| 39 | *R. cretica* | B. Özüdoğru 3439a (HUB) | Greece, Crete Island, around Impros, screes, 750 m, Latitude: 35.242861 N, Longitude 24.162797 E, 25.05.2013. | 3 | H37 / R33 |
| 40 | *R. cretica* | B. Özüdoğru 3439c (HUB) | Greece, Crete Island, Aradena valley, mobile screes, 580 m, Latitude: 35.220556 N, Longitude 24.058611 E, 25.05.2013. | 3 | H36 / R33, R34 |
| 41 | *R. cretica* | B. Özüdoğru 3439b (HUB) | Greece, Crete Island, 2-3 km from Anapoli Village to Aradena Valley, rocks, 613 m, Latitude: 35.222778 N, Longitude 24.072222 E, 25.05.2013. | 3 | H36 / R35, R36 |
| 42 | *R. cretica* | B. Özüdoğru 3440b (HUB) | Greece, Crete Island, Kortoluati Valley, screes, 196 m, Latitude: 35.197605 N, Longitude 24.465357 E, 26.05.2013. | 1 | H40 / R37 |
| 43 | *R. cretica* | B. Özüdoğru 3440a (HUB) | Greece, Crete Island, Illingos Valley, road side, screes, 65 m, Latitude: 35.205705 N, Longitude 24.125825 E, 26.05.2013. | 3 | H37 / R33, R38 |
| 44 | *R. cretica* | B. Özüdoğru CS2 (HUB) | Greece, Crete Island, around Kavusi Village, screes, 300 m, Latitude: 35.135 N, Longitude 25.866389 E, 27.05.2013. | 2 | H38, H39 / R39 |
| 45 | *R. cretica* | B. Özüdoğru CS3 (HUB) | Greece, Crete Island, opposite of Spinolongoa (Kalydon) island, mobile screes, 150 m, Latitude: 35.301386 N, Longitude 25.719923 E, 27.05.2013. | 2 | H41 / R39 |
| 46 | *R. varians* | H. Peşmen 2231 (HUB) | Turkey, Konya: Beyşehir, Kurucova, between Musalla and Muslu*, Pinus. nigra* and *Cedrus libani* forest, calcareous rocks, 1400-2000 m, 23.07.1975. | 1 | H68 / R40 |
| 47 | *R. varians* | B. Özüdoğru 3212 (HUB) | Turkey, Isparta: Aksu, Dedegöl Mountain, Kumçukuru, 2210 m, Latitude: 37.666667 N, Longitude 31.2575 E, 16. 07. 2011. | 1 | H70 / R40 |
| 48 | *R. varians* | B. Özüdoğru 3211 (HUB) | Turkey, Isparta: Aksu, Dedegöl Mountain, around Kuzukulağı plateau, southern sides, screes, 2200-2260 m, Latitude: 37.671944 N, Longitude 31.260556 E, 16.07.2011. | 2 | H70 / R40 |
| 49 | *R. varians* | B. Özüdoğru 3216 & K. Özgişi (HUB) | Turkey, Antalya: Akseki, Cevizli, Kuyucak village, Sivrikaya hill, mobile screes, 1680 m, Latitude: 37.2625 N, Longitude 31.2625 E, 22.07.2011. | 3 | H69 / R41 |
| 50 | *R. varians* | B. Özüdoğru 3524 & K. Özgişi (HUB) | Turkey, Antalya: Manavgat, Akpınar plateau, between Burunçalı and Kaldırım Mountain, screes, 1640 m, Latitude: 37.148439 N, Longitude 31.3255 E, 05.06.2014. | 3 | H67 / R42 |
| 51 | *R. lunaria* | A. Liston 7-82-29/1 (HUB) | Israel, coastal Galilee: opposite Shelomi, 27.02.1982. | 1 | H47 / R43 |
| 52 | *R. lunaria* | Michal Monosov 22250 (HUB) | Israel, Judean Mountains, Michal Monosov 22250, Latitude 31.800505 N, Longitude 35.239393 E, 19.05.2008, | 3 | H49 / R44, R45 |
| 53 | *R. lunaria* | Yair Ur 22721 (HUB) | Israel, Upper Jordan Valley, Yair Ur 22721, Latitude 32.903282, Longitude 35.554040, 10.04.2009. | 1 | H50 / R45 |
| 54 | *R. lunaria* | Ran Lotan 20466 (HUB) | Israel, South Golan, Ran Lotan 20466, Latitude 32.977777 N, Longitude 35.706818 E, 04, 09. 2007. | 1 | H48 / R46 |
| 55 | *R. sinuata* | A.A. Dönmez 13016 (HUB) | Turkey, Antalya: Alanya, Demittas¸ , Kocaoğlanlı village, Latitude 36.469333 N, Longitude 32.222928 E, macchie, 19.05.2006. | 1 | H60 / R47 |
| 56 | *R. sinuata* | B. Özüdoğru 3540 & K. Özgişi (HUB) | Turkey, Antalya: Alanya, Demittas¸ , Kocaoğlanlı village, macchie, 300 m, 300 m Latitude 36.469333 N, Longitude 32.222928 E, 05.06.2014. | 2 | H61 / R47, R48 |
| 57 | *R. sinuata* | B. Özüdoğru 2882 (HUB) | Turkey, Antalya: between İbradı and Manavgat, around Yaylaalan Village, macchie, 480 m, Latitude 36.944836 N, Longitude 31.500916 E, 29.04.2011. | 3 | H63, H64 / R49 |
| 58 | *R. sinuata* | B. Özüdoğru 3436 & G.Akaydın (HUB) | Turkey, Antalya: Alanya, Hacımehmetli village, road side, on rocks, 113 m, Latitude 36.56278 N, Longitude 31.96056 E, 18.05.2013, | 3 | H51, H52 / R50 |
| 59 | *R. sinuata* | B. Özüdoğru 3435 & G. Akaydın (HUB) | Turkey, Antalya: Akseki, around Fersin Village, rocks, 490 m, Latitude 36.795 N, Longitude 31.768056 E, 18.05.2013. | 4 | H62 / R51 |
| 60 | *R. sinuata* | B. Özüdoğru 3347 (HUB) | Turkey, Antalya: Manavgat, above Ahmetler village, under *P. brutia*, mobile screes, 880–900 m, Latitude 36.848127 N, Longitude 31.716949 E, 05.06.2012. | 2 | H62 / R50, R51 |
| 61 | *R. sinuata* | B. Özüdoğru 3501 & E. Çilden (HUB) | Turkey, Antalya: Gündoğmuş, around Narağacı Village, mobile screes, 460-490 m, Latitude 36.782756 N, Longitude 32.103364 E, 23.05.2014. | 3 | H57 / R50 |
| 62 | *R. sinuata* | B. Özüdoğru 2947 & D. Töre (HUB) | Turkey, Hatay: Samandağ, Çevlik, road sides, 17 m, Latitude 36.149508 N, Longitude 35.903146 E, 13.05.2011. | 3 | H54, H55 / R52, R53 |
| 63 | *R. sinuata* | B. Özüdoğru 2953 & D. Töre (HUB) | Turkey, Hatay: Belen, between Kömürçukuru and Tahtaköprü villages, 350–450 m, mobile screes, Latitude 36.391353 N, Longitude 36.161059 E, 14.05.2011. | 4 | H53 / R54 |
| 64 | *R. sinuata* | H. Altınözlü 2012-1 (HUB) | Turkey, Adana: between Feke and Sülemişli, roadside, 620 m, , Latitude 37.90639 N, Longitude 35.94528 E, 16.04.2012. | 3 | H56 / R55 |
| 65 | *R. sinuata* | B. Özüdoğru 3317 & L. Tutar (HUB) | Turkey, Kahramanmaraş: between Andırın and Kadirli, Harboğazı, mobile screes, 260 m, Latitude 37.466204 N, Longitude 36.341106 E, 11.05.2012. | 5 | H58, H59 / R56 |
