## Supplementary material for "Ecological specialization promotes diversity and diversification in the East Mediterranean genus *Ricotia* (Brassicaceae)": Suppementary Information: Table S2.docx

**Table S2.** First-order texture measures

| **Metric** | **Measure** | **Value Range** | **Expected Relationship with Heterogeneity** |
| --- | --- | --- | --- |
| Coefficient of variation | Normalized dispersion of EVI | >=0 | Positive |
| Evenness | Evenness of EVI | >=0; <=1 | Positive |
| Range | Range of EVI | >=0 | Positive |
| Shannon | Diversity of EVI | >=0; <=ln(max # of different EVI) | Positive |
| Simpson | Diversity of EVI | >=0; <=1-1/(max # of different EVI) | Positive |
| Standard deviation | Dispersion of EVI | >=0 | Positive |
