## Supplementary material for "Ecological specialization promotes diversity and diversification in the East Mediterranean genus *Ricotia* (Brassicaceae)": Suppementary Information: Table S3.docx

**Table S3.** Second-order texture measures

| **Metric** | **Measure** | **Value Range** | **Expected Relationship with Heterogeneity** |
| --- | --- | --- | --- |
| Contrast | Exponentially weighted difference in EVI between adjacent pixels | >=0 | Positive |
| Correlation | Linear dependency of EVI on adjacent pixels | >=-1; <=1 | Nonlinear |
| Dissimilarity | Difference in EVI between adjacent pixels | >=0 | Positive |
| Entropy | Disorderliness of EVI | >=0 | Positive |
| Homogeneity | Similarity of EVI between adjacent pixels | >=0; <=1 | Negative |
| Maximum | Dominance of EVI combinations between adjacent pixels | >=0; <=1 | Negative |
| Uniformity | Orderliness of EVI | >=0; <=1 | Negative |
| Variance | Dispersion of EVI combinations between adjacent pixels | >=0 | Positive |
