## Supplementary material for "Ecological specialization promotes diversity and diversification in the East Mediterranean genus *Ricotia* (Brassicaceae)": Suppementary Information: Table S4.docx

**Table S4.** Defined ITS ribotypes in this study with their voucher and Genbank accession numbersvoucher numbers.

| Ribotype | n | Species | Voucher | Genebank numbers |
| --- | --- | --- | --- | --- |
| R1 | 3 | *R. aucheri* | B. Özüdoğru 3561 & E.Çilden (HUB) | MK875183 |
| R2 | 9 | *R. aucheri* | B. Özüdoğru 3350 (HUB), B. Özüdoğru 3352 (HUB), B. Özüdoğru 3353 (HUB) | MK875184 |
| R3 | 1 | *R. aucheri* | B. Özüdoğru 3353 (HUB) | MK875185 |
| R4 | 7 | *R. aucheri* | MYP Tunceli-t (HUB), H. Yıldırım 3101 (HUB) | MK875186 |
| R5 | 1 | *R. aucheri* | Ş. Yıldırımlı 2298 (HUB) | MK875187 |
| R6 | 5 | *R. aucheri* | A. Güner 4760 (HUB), B. Özüdoğru 3562 (HUB), B. Özüdoğru 3565 (HUB) | MK875188 |
| R7 | 4 | *R. aucheri* | A. Tatlı 4255 (HUB), B. Özüdoğru 3563 (HUB) | MK875189 |
| R8 | 2 | *R. aucheri* | B. Özüdoğru 3569 (HUB) | MK875190 |
| R9 | 3 | *R. aucheri* | B. Özüdoğru 3570 (HUB) | MK875191 |
| R10 | 3 | *R. aucheri* | B. Özüdoğru 3565 (HUB), H. Yıldırım 3120 (HUB) | MK875192 |
| R11 | 1 | *R. aucheri* | G. Zare (HUB) | MK875193 |
| R12 | 2 | *R. aucheri* | B. Özüdoğru 3471 & G. Akaydın (HUB) | MK875194 |
| R13 | 1 | *R. aucheri* | B. Özüdoğru 3471 & G. Akaydın (HUB) | MK875195 |
| R14 | 3 | *R. aucheri* | B.Özüdoğru 3472 & G.Akaydın (HUB) | MK875196 |
| R15 | 3 | *R. aucheri* | B. Özüdoğru 3217 & K. Özgişi (HUB) | MK875197 |
| R16 | 1 | *R. aucheri* | A. Güner 3638 (HUB) | MK875198 |
| R17 | 1 | *R. aucheri* | H. Yıldırım 2723 (HUB) | MK875199 |
| R18 | 2 | *R. aucheri* | B. Özüdoğru 3354 (HUB) | MK875200 |
| R19 | 1 | *R. aucheri* | B. Özüdoğru 3354 (HUB) | MK875201 |
| R20 | 4 | *R. aucheri* | B. Özüdoğru 3354 (HUB), B. Özüdoğru 3559 & E. Çilden (HUB) | MK875202 |
| R21 | 3 | *R. aucheri* | B. Özüdoğru 3545 & E. Çilden (HUB) | MK875203 |
| R22 | 2 | *R.isatoides* | B. Özüdoğru 3475 (HUB) | MK875204 |
| R23 | 3 | *R.tenuifolia* | B. Özüdoğru 2968 (HUB), Paksoy 1080 (HUB) | MK875205 |
| R24 | 1 | *R.tenuifolia* | B. Özüdoğru 2968 (HUB) | MK875206 |
| R25 | 4 | *R.davisiana* | B. Özüdoğru 3215 & K.Özgişi (HUB), H. Peşmen 4619 & A. Güner (HUB) | MK875207 |
| R26 | 9 | *R.carnosula* | B. Özüdoğru 2852 (HUB), B. Özüdoğru 2862 (HUB), B. Özüdoğru 2863 (HUB), B. Özüdoğru 3437 & G. Akaydın (HUB), B. Özüdoğru, 2857 (HUB) | MK875208 |
| R27 | 2 | *R.carnosula* | B. Özüdoğru 2883 (HUB), B. Özüdoğru 2884 (HUB) | MK875209 |
| R28 | 1 | *R.carnosula* | B. Özüdoğru 2884 (HUB) | MK875210 |
| R29 | 1 | *R.carnosula* | B. Özüdoğru 2863 (HUB) | MK875211 |
| R30 | 2 | *R.carnosula* | B. Özüdoğru 3315 (HUB) | MK875212 |
| R31 | 2 | *R.carnosula* | B. Özüdoğru 3315 (HUB) | MK875213 |
| R32 | 3 | *R.carnosula* | B. Özüdoğru 3316 (HUB) | MK875214 |
| R33 | 6 | *R.cretica* | B. Özüdoğru 3439a (HUB), B. Özüdoğru 3439c (HUB), B. Özüdoğru 3440a (HUB) | MK875215 |
| R34 | 2 | *R.cretica* | B. Özüdoğru 3439c (HUB) | MK875216 |
| R35 | 2 | *R.cretica* | B. Özüdoğru 3439b (HUB) | MK875217 |
| R36 | 1 | *R.cretica* | B. Özüdoğru 3439b (HUB) | MK875218 |
| R37 | 1 | *R.cretica* | B. Özüdoğru 3440b (HUB) | MK875219 |
| R38 | 1 | *R.cretica* | B. Özüdoğru 3440a (HUB) | MK875220 |
| R39 | 4 | *R.cretica* | B. Özüdoğru CS2 (HUB), B. Özüdoğru CS3 (HUB) | MK875221 |
| R40 | 4 | *R.varians* | H. Peşmen 2231 (HUB), B. Özüdoğru 3212 (HUB), B. Özüdoğru 3211 (HUB) | MK875222 |
| R41 | 3 | *R.varians* | B. Özüdoğru 3216 & K. Özgişi (HUB) | MK875223 |
| R42 | 3 | *R.varians* | B. Özüdoğru 3524 & K. Özgişi (HUB) | MK875224 |
| R43 | 1 | *R.lunaria* | A. Liston 7-82-29/1 (HUB) | MK875225 |
| R44 | 2 | *R.lunaria* | Michal Monosov 22250 (HUB) | MK875226 |
| R45 | 2 | *R.lunaria* | Michal Monosov 22250 (HUB), Yair Ur 22721 (HUB) | MK875227 |
| R46 | 1 | *R.lunaria* | Ran Lotan 20466 (HUB) | MK875228 |
| R47 | 2 | *R.sinuata* | A.A. Dönmez 13016 (HUB), B. Özüdoğru 3540 & K. Özgişi (HUB) | MK875229 |
| R48 | 1 | *R.sinuata* | B. Özüdoğru 3540 & K. Özgişi (HUB) | MK875230 |
| R49 | 3 | *R.sinuata* | B. Özüdoğru 2882 (HUB) | MK875231 |
| R50 | 7 | *R.sinuata* | B. Özüdoğru 3436 & G.Akaydın, B. Özüdoğru 3347 (HUB), B. Özüdoğru 3501 & E. Çilden (HUB) | MK875232 |
| R51 | 5 | *R.sinuata* | B. Özüdoğru 3435 & G. Akaydın (HUB), B. Özüdoğru 3347 (HUB) | MK875233 |
| R52 | 2 | *R.sinuata* | B. Özüdoğru 2947 & D. Töre (HUB) | MK875234 |
| R53 | 1 | *R.sinuata* | B. Özüdoğru 2947 & D. Töre (HUB) | MK875235 |
| R54 | 4 | *R.sinuata* | B. Özüdoğru 2953 & D. Töre (HUB) | MK875236 |
| R55 | 3 | *R.sinuata* | H. Altınözlü 2012-1 (HUB) | MK875237 |
| R56 | 5 | *R.sinuata* | B. Özüdoğru 3317 & L. Tutar (HUB) | MK875238 |
