## Supplementary material for "Ecological specialization promotes diversity and diversification in the East Mediterranean genus *Ricotia* (Brassicaceae)": Suppementary Information: Table S5.docx

**Table S5.** Defined cpDNA haplotypes in this study with their voucher and Genbank accession numbers

| Haplotype | n | Species | Voucher | Genebank numbers *trn*L-F / *trn*Q-5’ rps16 |
| --- | --- | --- | --- | --- |
| H1 | 1 | *R.aucheri* | A. Güner 3638 (HUB) | MK941911 / MK941981 |
| H2 | 11 | *R.aucheri* | A. Güner 4760 (HUB), A. Tatlı 4255 (HUB), G. Zare (HUB), B. Özüdoğru 3562 (HUB), B. Özüdoğru 3563 (HUB), B. Özüdoğru 3565 (HUB) | MK941912 / MK941982 |
| H3 | 3 | *R.aucheri* | B. Özüdoğru, 3353 (HUB) | MK941913 / MK941983 |
| H4 | 1 | *R.aucheri* | B. Özüdoğru, 3353 (HUB) | MK941914 / MK941984 |
| H5 | 1 | *R.aucheri* | B. Özüdoğru 3471 & G. Akaydın (HUB) | MK941915 / MK941985 |
| H6 | 2 | *R.aucheri* | B. Özüdoğru 3471 & G. Akaydın (HUB) | MK941916 / MK941986 |
| H7 | 1 | *R.aucheri* | B. Özüdoğru 3570 (HUB) | MK941917 / MK941987 |
| H8 | 2 | *R.aucheri* | B. Özüdoğru 3570 (HUB) | MK941918 / MK941988 |
| H9 | 3 | *R.aucheri* | B. Özüdoğru 3354 (HUB) | MK941919 / MK941989 |
| H10 | 1 | *R.aucheri* | B. Özüdoğru 3354 (HUB) | MK941920 / MK941990 |
| H11 | 3 | *R.aucheri* | B. Özüdoğru 3217 & K. Özgişi (HUB) | MK941921 / MK941991 |
| H12 | 3 | *R.aucheri* | B. Özüdoğru 3559 E. Çilden (HUB) | MK941922 / MK941992 |
| H13 | 1 | *R.aucheri* | H. Yıldırım 2723 (HUB) | MK941923 / MK941993 |
| H14 | 3 | *R.aucheri* | H. Yıldırım 3101 (HUB) | MK941924 / MK941994 |
| H15 | 1 | *R.aucheri* | H. Yıldırım 3101 (HUB) | MK941925 / MK941995 |
| H16 | 1 | *R.aucheri* | H. Yıldırım 3120 (HUB) | MK941926 / MK941996 |
| H17 | 1 | *R.aucheri* | B. Özüdoğru 3562 (HUB) | MK941927 / MK941997 |
| H18 | 3 | *R.aucheri* | B. Özüdoğru 3545 & E. Çilden (HUB) | MK941928 / MK941998 |
| H19 | 2 | *R.aucheri* | MYP Tunceli-t (HUB) | MK941929 / MK941999 |
| H20 | 1 | *R.aucheri* | MYP Tunceli-t (HUB) | MK941930 / MK942000 |
| H21 | 3 | *R.aucheri* | B. Özüdoğru 3561 & E.Çilden (HUB) | MK941931 / MK942001 |
| H22 | 1 | *R.aucheri* | B.Özüdoğru 3472 & G.Akaydın (HUB) | MK941932 / MK942002 |
| H23 | 2 | *R.aucheri* | B.Özüdoğru 3472 & G.Akaydın (HUB) | MK941933 / MK942003 |
| H24 | 6 | *R.aucheri* | B. Özüdoğru 3350 (HUB), B. Özüdoğru 3352 (HUB) | MK941934 / MK942004 |
| H25 | 2 | *R.aucheri* | B. Özüdoğru 3569 (HUB) | MK941935 / MK942005 |
| H26 | 1 | *R.aucheri* | Ş. Yıldırımlı 2298 (HUB) | MK941936 / MK942006 |
| H27 | 4 | *R.carnosula* | B. Özüdoğru 2884 (HUB), B. Özüdoğru 2863 (HUB), B. Özüdoğru 3437 & G. Akaydın (HUB) | MK941937 / MK942007 |
| H28 | 4 | *R.carnosula* | B. Özüdoğru 2852 (HUB), B. Özüdoğru, 2857 (HUB) | MK941938 / MK942008 |
| H29 | 1 | *R.carnosula* | B. Özüdoğru 2863 (HUB) | MK941939 / MK942009 |
| H30 | 3 | *R.carnosula* | B. Özüdoğru 3316 (HUB) | MK941940 / MK942010 |
| H31 | 4 | *R.carnosula* | B. Özüdoğru 3315 (HUB) | MK941941 / MK942011 |
| H32 | 1 | *R.carnosula* | B. Özüdoğru 2883 (HUB) | MK941942 / MK942012 |
| H33 | 1 | *R.carnosula* | B. Özüdoğru 3437 & G. Akaydın (HUB) | MK941943 / MK942013 |
| H34 | 1 | *R.carnosula* | B. Özüdoğru 2862 (HUB) | MK941944 / MK942014 |
| H35 | 1 | *R.carnosula* | B. Özüdoğru 2862 (HUB) | MK941945 / MK942015 |
| H36 | 6 | *R.cretica* | B. Özüdoğru 3439b (HUB), B. Özüdoğru 3439c (HUB) | MK941946 / MK942016 |
| H37 | 6 | *R.cretica* | B. Özüdoğru 3440a (HUB), B. Özüdoğru 3439a (HUB) | MK941947 / MK942017 |
| H38 | 1 | *R.cretica* | B. Özüdoğru CS2 (HUB) | MK941948 / MK942018 |
| H39 | 1 | *R.cretica* | B. Özüdoğru CS2 (HUB) | MK941949 / MK942019 |
| H40 | 1 | *R.cretica* | B. Özüdoğru 3440b (HUB) | MK941950 / MK942020 |
| H41 | 2 | *R.cretica* | B. Özüdoğru CS3 (HUB) | MK941951 / MK942021 |
| H42 | 1 | *R.davisiana* | H. Peşmen 4619 & A. Güner (HUB) | MK941952 / MK942022 |
| H43 | 2 | *R.davisiana* | B. Özüdoğru 3215 & K.Özgişi (HUB) | MK941953 / MK942023 |
| H44 | 1 | *R.davisiana* | B. Özüdoğru 3215 & K.Özgişi (HUB) | MK941954 / MK942024 |
| H45 | 1 | *R.isatoides* | B. Özüdoğru 3475 (HUB) | MK941955 / MK942025 |
| H46 | 1 | *R.isatoides* | B. Özüdoğru 3475 (HUB) | MK941956 / MK942026 |
| H47 | 1 | *R.lunaria* | A. Liston 7-82-29/1 (HUB) | MK941957 / MK942027 |
| H48 | 1 | *R.lunaria* | Ran Lotan 20466 (HUB) | MK941958 / MK942028 |
| H49 | 3 | *R.lunaria* | Michal Monosov 22250 (HUB) | MK941959 / MK942029 |
| H50 | 1 | *R.lunaria* | Yair Ur 22721 (HUB) | MK941960 / MK942030 |
| H51 | 2 | *R.sinuata* | B. Özüdoğru 3436 & G.Akaydın | MK941961 / MK942031 |
| H52 | 1 | *R.sinuata* | B. Özüdoğru 3436 & G.Akaydın | MK941962 / MK942032 |
| H53 | 4 | *R.sinuata* | B. Özüdoğru 2953 & D. Töre (HUB) | MK941963 / MK942033 |
| H54 | 2 | *R.sinuata* | B. Özüdoğru 2947 & D. Töre (HUB) | MK941964 / MK942034 |
| H55 | 1 | *R.sinuata* | B. Özüdoğru 2947 & D. Töre (HUB) | MK941965 / MK942035 |
| H56 | 3 | *R.sinuata* | H. Altınözlü 2012-1 (HUB) | MK941966 / MK942036 |
| H57 | 3 | *R.sinuata* | B. Özüdoğru 3501 & E. Çilden (HUB) | MK941967 / MK942037 |
| H58 | 4 | *R.sinuata* | B. Özüdoğru 3317 & L. Tutar (HUB) | MK941968 / MK942038 |
| H59 | 1 | *R.sinuata* | B. Özüdoğru 3317 & L. Tutar (HUB) | MK941969 / MK942039 |
| H60 | 1 | *R.sinuata* | A.A. Dönmez 13016 (HUB) | MK941970 / MK942040 |
| H61 | 2 | *R.sinuata* | B. Özüdoğru 3540 & K. Özgişi (HUB) | MK941971 / MK942041 |
| H62 | 6 | *R.sinuata* | B. Özüdoğru 3435 & G. Akaydın (HUB), B. Özüdoğru 3347 (HUB) | MK941972 / MK942042 |
| H63 | 2 | *R.sinuata* | B. Özüdoğru 2882 (HUB) | MK941973 / MK942043 |
| H64 | 1 | *R.sinuata* | B. Özüdoğru 2882 (HUB) | MK941974 / MK942044 |
| H65 | 3 | *R.tenuifolia* | B. Özüdoğru 2968 (HUB) | MK941975 / MK942045 |
| H66 | 1 | *R.tenuifolia* | Paksoy 1080 (HUB) | MK941976 / MK942046 |
| H67 | 3 | *R.varians* | B. Özüdoğru 3524 & K. Özgişi (HUB) | MK941977 / MK942047 |
| H68 | 1 | *R.varians* | H. Peşmen 2231 (HUB) | MK941978 / MK942048 |
| H69 | 3 | *R.varians* | B. Özüdoğru 3216 & K. Özgişi (HUB) | MK941979 / MK942049 |
| H70 | 3 | *R.varians* | B. Özüdoğru 3211 (HUB), B. Özüdoğru 3212 (HUB) | MK941980 / MK942050 |
