## Supplementary material for "Ecological specialization promotes diversity and diversification in the East Mediterranean genus *Ricotia* (Brassicaceae)": Suppementary Information: Table S6.docx

|  | N | SL | S | H | Hd | π | D | | Fs |
| --- | --- | --- | --- | --- | --- | --- | --- | --- | --- |
| ***Ricotia*** | 156 | 599 | (127)155 | 56 | 0.9798 | 0.05561 | 0.56004 | | 1.298 |
| ***Tenuifolia* clade** | 6 | 579 | (14)15 | 3 | 0.733 | 0.01257 | 1.13697 | | 3.922 |
| **Core clade** | 90 | 599 | (66)68 | 32 | 0.9650 | 0.02227 | -0.07471 | | -1.604 |
| ***R. aucheri*** | 60 | 592 | 51 | 21 | 0.944 | 0.01256 | -0.88156 | | -1.978 |
| ***R. carnosula*** | 20 | 598 | 9 | 7 | 0.779 | 0.00239 | -1.50541 | | -2.281 |
| ***R. cretica*** | 17 | 598 | 10 | 7 | 0.831 | 0.00586 | 0.67103 | 0.070 | |
| ***R. sinuata*** | 33 | 599 | 18 | 10 | 0.896 | 0.00715 | 0.12384 | 0.104 | |
| ***R.varians*** | 10 | 598 | 6 | 3 | NA | NA | NA | NA | |
| ***R. lunaria*** | 6 | 598 | 6 | 4 | NA | NA | NA | NA | |
| ***R. tenuifolia*** | 4 | NA | NA | NA | NA | NA | NA | NA | |
| ***R. isatoides*** | 4 | NA | NA | NA | NA | NA | NA | NA | |
| ***R. davisiana*** | 4 | NA | NA | NA | NA | NA | NA | NA | |

**Table S4**. Summary of genetic diversity indices and results of neutrality tests (Tajima’s D and Fu’s Fs) for nuclear ITS data. N, number of sequences; SL, sequence length (bp); S, number of segregating sites with alignment gaps excluded (in parenthesis) or included; H, number of haplotypes; Hd, haplotype diversity; π, nucleotide diversity.
