## Supplementary material for "Ecological specialization promotes diversity and diversification in the East Mediterranean genus *Ricotia* (Brassicaceae)": Suppementary Information: Table S7.docx

**Table S5.** Summary of genetic diversity indices (Θs and p) and results of neutrality tests (Tajima’s D and Fu’s Fs) for combined chloroplast (*trn*L-F + *trn*Q-*rps*16) data. N, number of sequences; SL, sequence length (bp); S, number of segregating sites with alignment gaps excluded (in parenthesis) or included; H, number of haplotypes; Hd, haplotype diversity; π, nucleotide diversity.

| Species | N | SL | S | nH | Hd | π | D | | Fs |
| --- | --- | --- | --- | --- | --- | --- | --- | --- | --- |
| ***Ricotia*** | 156 | 1244 | (138)484 | 70 | 0.983 | 0.02897 | -0.23796 | | -3.203 |
| ***Tenuifolia* clade** | 6 | 1202 | (25)62 | 4 | 0.800 | 0.01167 | 1.22593 | | 5.771 |
| **Core clade** | 90 | 1225 | (72)346 | 40 | 0.975 | 0.01643 | 0.09533 | | -1.737 |
| ***R. aucheri*** | 60 | 1201 | 76 | 26 | 0.945 | 0.01690 | 0.20433 | | 1.027 |
| ***R. carnosula*** | 20 | 1232 | 36 | 9 | 0.889 | 0.00767 | -0.37606 | | 1.053 |
| ***R. carnosula*!** | 17 | 1232 | 7 | 8 | 0.868 | 0.00136 | -0.65756 | | -2.121 |
| ***R. cretica*** | 17 | 1229 | 16 | 6 | 0.772 | 0.00426 | 0.41171 | | 2.325 |
| ***R. sinuata*** | 33 | 1225 | 21 | 12 | 0.922 | 0.00603 | 0.48345 | | 1.377 |
| ***R. lunaria*** | 6 | NA | NA | NA | NA | NA | NA | | NA |
| ***R. varians*** | 10 | NA | NA | NA | NA | NA | NA | NA | |
| ***R. tenuifolia*** | 4 | NA | NA | NA | NA | NA | NA | NA | |
| ***R. isatoides*** | 4 | NA | NA | NA | NA | NA | NA | NA | |
| ***R. davisiana*** | 4 | NA | NA | NA | NA | NA | NA | NA | |
