## Supplementary material for "Ecological specialization promotes diversity and diversification in the East Mediterranean genus *Ricotia* (Brassicaceae)": Suppementary Information: Table S8.docx

**Table S8.** Results of the BioGeoBEARS analysis of the cp dataset. The DEC+J model (in **boldface**) was recovered as fitting the data significantly better than the five other models.

| Model | LnL | Number of parameters | Parameters | | |
| --- | --- | --- | --- | --- | --- |
|  |  |  | d | e | j |
| DEC | -42.81 | 2 | 0.0044 | 1.6e-09 | 0 |
| DEC+J | **-39.59** | **3** | **0.0004** | **1.0e-12** | **0.0090** |
| DIVALIKE | -44.18 | 2 | 0.0065 | 1.0e-12 | 0 |
| DIVALIKE+J | -41.04 | 3 | 0.0011 | 1.0e-12 | 0.010 |
| BAYAREALIKE | -60.79 | 2 | 0.0061 | 0.047 | 0 |
| BAYAREALIKE+J | -41.74 | 3 | 1.0e-07 | 1.0e-07 | 0.012 |
