## Supplementary material for "Ecological specialization promotes diversity and diversification in the East Mediterranean genus *Ricotia* (Brassicaceae)": Suppementary Information: Table S9.docx

**Table S9** List of three most possible character states at nodes, with their corresponding probability (p) as returned by the DEC+J analysis implemented in BioGeoBEARS. The most likely distribution prior speciation is indicated in boldface. Node numbers refers to nodes shown in supplementary Figure S8.

| Node numbers | Character state | Probability | Character state | Probability | Character state | Probability |
| --- | --- | --- | --- | --- | --- | --- |
| 1 | **D** | 100.000000 | BD | 0.000000 | DF | 0.000000 |
| 2 | **D** | 100.000000 | BD | 0.000000 | DF | 0.000000 |
| 3 | **D** | 100.000000 | BD | 0.000000 | DF | 0.000000 |
| 4 | **D** | 100.000000 | BD | 0.000000 | DF | 0.000000 |
| 5 | **D** | 100.000000 | BD | 0.000000 | DF | 0.000000 |
| 6 | **D** | 100.000000 | BD | 0.000000 | DF | 0.000000 |
| 7 | **D** | 100.000000 | BD | 0.000000 | DF | 0.000000 |
| 8 | **D** | 100.000000 | BD | 0.000000 | DF | 0.000000 |
| 9 | **D** | 100.000000 | BD | 0.000000 | DF | 0.000000 |
| 10 | **D** | 100.000000 | BD | 0.000000 | DF | 0.000000 |
| 11 | **D** | 100.000000 | BD | 0.000000 | DF | 0.000000 |
| 12 | **D** | 100.000000 | BD | 0.000000 | DF | 0.000000 |
| 13 | **D** | 100.000000 | BD | 0.000000 | DF | 0.000000 |
| 14 | **D** | 100.000000 | BD | 0.000000 | DF | 0.000000 |
| 15 | **D** | 100.000000 | BD | 0.000000 | DF | 0.000000 |
| 16 | **D** | 100.000000 | BD | 0.000000 | DF | 0.000000 |
| 17 | **D** | 100.000000 | BD | 0.000000 | DF | 0.000000 |
| 18 | **D** | 100.000000 | BD | 0.000000 | DF | 0.000000 |
| 19 | **D** | 100.000000 | BD | 0.000000 | DF | 0.000000 |
| 20 | **D** | 100.000000 | BD | 0.000000 | DF | 0.000000 |
| 21 | **D** | 100.000000 | BD | 0.000000 | DF | 0.000000 |
| 22 | **D** | 100.000000 | BD | 0.000000 | DF | 0.000000 |
| 23 | **D** | 100.000000 | BD | 0.000000 | DF | 0.000000 |
| 24 | **D** | 100.000000 | BD | 0.000000 | DF | 0.000000 |
| 25 | **D** | 100.000000 | BD | 0.000000 | DF | 0.000000 |
| 26 | **B** | 100.000000 | AB | 0.000000 | BC | 0.000000 |
| 27 | B | 100.000000 | AB | 0.000000 | BC | 0.000000 |
| 28 | **B** | 100.000000 | AB | 0.000000 | BC | 0.000000 |
| 29 | **B** | 100.000000 | AB | 0.000000 | BC | 0.000000 |
| 30 | **B** | 100.000000 | AB | 0.000000 | BC | 0.000000 |
| 31 | **B** | 100.000000 | AB | 0.000000 | BC | 0.000000 |
| 32 | **B** | 100.000000 | AB | 0.000000 | BC | 0.000000 |
| 33 | **A** | 100.000000 | AB | 0.000000 | AC | 0.000000 |
| 34 | **A** | 100.000000 | AB | 0.000000 | AC | 0.000000 |
| 35 | **A** | 100.000000 | AB | 0.000000 | AC | 0.000000 |
| 36 | **A** | 100.000000 | AB | 0.000000 | AC | 0.000000 |
| 37 | **A** | 100.000000 | AB | 0.000000 | B | 0.0000000 |
| 38 | **B** | 93.967500 | AB | 5.059509 | A | 0.972992 |
| 39 | **B** | 100.000000 | BC | 0.000000 | AB | 0.000000 |
| 40 | **B** | 100.000000 | BC | 0.000000 | AB | 0.000000 |
| 41 | **B** | 100.000000 | BC | 0.000000 | AB | 0.000000 |
| 42 | **B** | 99.444570 | AB | 0.448715 | A | 0.106710 |
| 43 | **E** | 100.000000 | CE | 0.000000 | BE | 0.000000 |
| 44 | **E** | 100.000000 | CE | 0.000000 | BE | 0.000000 |
| 45 | **E** | 100.000000 | CE | 0.000000 | BE | 0.000000 |
| 46 | **C** | 100.000000 | CE | 0.000000 | BC | 0.000000 |
| 47 | **C** | 100.000000 | CE | 0.000000 | BC | 0.000000 |
| 48 | **C** | 75.339020 | E | 23.410530 | CE | 1.250448 |
| 49 | **B** | 100.000000 | BC | 0.000000 | BE | 0.000000 |
| 50 | **B** | 100.000000 | BC | 0.000000 | BE | 0.000000 |
| 51 | **B** | 100.000000 | BC | 0.000000 | BE | 0.000000 |
| 52 | **B** | 100.000000 | BC | 0.000000 | BE | 0.000000 |
| 53 | **B** | 100.000000 | BC | 0.000000 | BE | 0.000000 |
| 54 | **B** | 100.000000 | BC | 0.000000 | BE | 0.000000 |
| 55 | **B** | 100.000000 | BC | 0.000000 | BE | 0.000000 |
| 56 | **B** | 100.000000 | BC | 0.000000 | BE | 0.000000 |
| 57 | **B** | 100.000000 | BC | 0.000000 | BE | 0.000000 |
| 58 | **B** | 100.000000 | BC | 0.000000 | BE | 0.000000 |
| 59 | **B** | 100.000000 | BC | 0.000000 | BE | 0.000000 |
| 60 | **C** | 100.000000 | BC | 0.000000 | CE | 0.000000 |
| 61 | **C** | 100.000000 | BC | 0.000000 | CE | 0.000000 |
| 62 | **B** | 45.613950 | BC | 30.738490 | C | 23.647560 |
| 63 | **B** | 42.281250 | BC | 32.055100 | C | 23.186150 |
| 64 | **B** | 93.643200 | BC | 5.304700 | C | 0.459293 |
| 65 | **A** | 100.000000 | AB | 0.000000 | AD | 0.000000 |
| 66 | **B** | 100.000000 | AB | 0.000000 | BD | 0.000000 |
| 67 | **B** | 90.126700 | AB | 8.327513 | A | 1.545791 |
| 68 | **B** | 98.281720 | A | 0.657053 | BC | 0.416079 |
| 69 | **B** | 75.724720 | D | 21.197350 | BD | 2.361748 |
