## Supplementary material for "Ecological specialization promotes diversity and diversification in the East Mediterranean genus *Ricotia* (Brassicaceae)": Suppementary Information: Table S10.docx

**Table S10.** Percent variable contributions according to permutation importance in Ecological Niche Modelling (ENM) analyses. For variable abbreviation see Appendix S2.

| ***R. aucheri*** | | ***R. cretica*** | | ***R. carnosula*** | | ***R. lunaria*** | | ***R. sinuata*** | |
| --- | --- | --- | --- | --- | --- | --- | --- | --- | --- |
| Variable | Permutation importance | Variable | Permutation importance | Variable | Permutation importance | Variable | Permutation importance | Variable | Permutation importance |
| cv | 31.2 | slope | 29.7 | taxnwrb | 53.3 | bio15 | 63.3 | bio12 | 84.9 |
| bio4 | 21 | bio15 | 25.2 | bio1 | 17.3 | bio14 | 25.2 | bio15 | 4.3 |
| bio12 | 18.2 | bio1 | 11.3 | bio12 | 11.2 | bio12 | 6.8 | bio14 | 3.2 |
| bio15 | 11.6 | cv | 10.4 | bio3 | 6.8 | cv | 1.7 | slope | 2.3 |
| contrast | 11 | bio2 | 9.9 | aspect | 5.7 | contrast | 0.8 | cv | 1.6 |
| bio1 | 2.4 | aspect | 4.9 | bio2 | 2.4 | bio5 | 0.5 | bio1 | 0.9 |
| correlation | 1.5 | evenness | 3.9 | slope | 1.7 | bio4 | 0.4 | bio4 | 0.6 |
| slope_tr | 1.2 | bio12 | 1.9 | entropy | 0.5 | bio2 | 0.4 | bio3 | 0.6 |
| bio2 | 0.9 | correlation | 1.1 | correlation | 0.4 | slope | 0.4 | contrast | 0.6 |
| bio14 | 0.3 | entropy | 0.8 | contrast | 0.4 | bio3 | 0.1 | aspect | 0.4 |
| bio8 | 0.3 | homogeneity | 0.5 | evenness | 0.2 | aspect | 0.1 | bio2 | 0.3 |
| taxnwrbr | 0.2 | taxnwrb | 0.4 | cv | 0.1 | entropy | 0.1 | evenness | 0.2 |
| bio3 | 0.1 | contrast | 0 |  |  | correlation | 0.1 | taxnwrb | 0.1 |
