## Supplementary figures and images for "Ecological specialization promotes diversity and diversification in the East Mediterranean genus *Ricotia* (Brassicaceae)"

### Figure S2.pdf

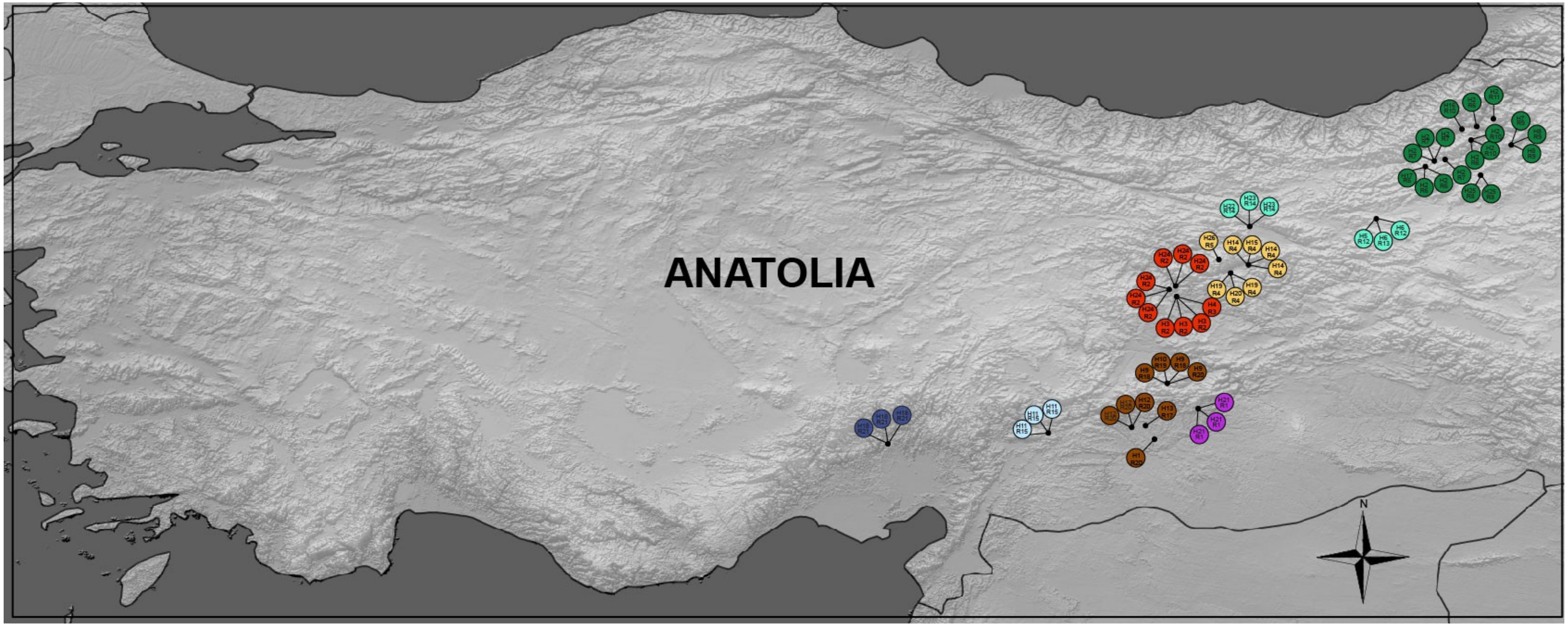


**Figure S2.** Spatial distribution of haplotypes and ribotypes in *Ricotia aucheri*.

### Figure S3.pdf

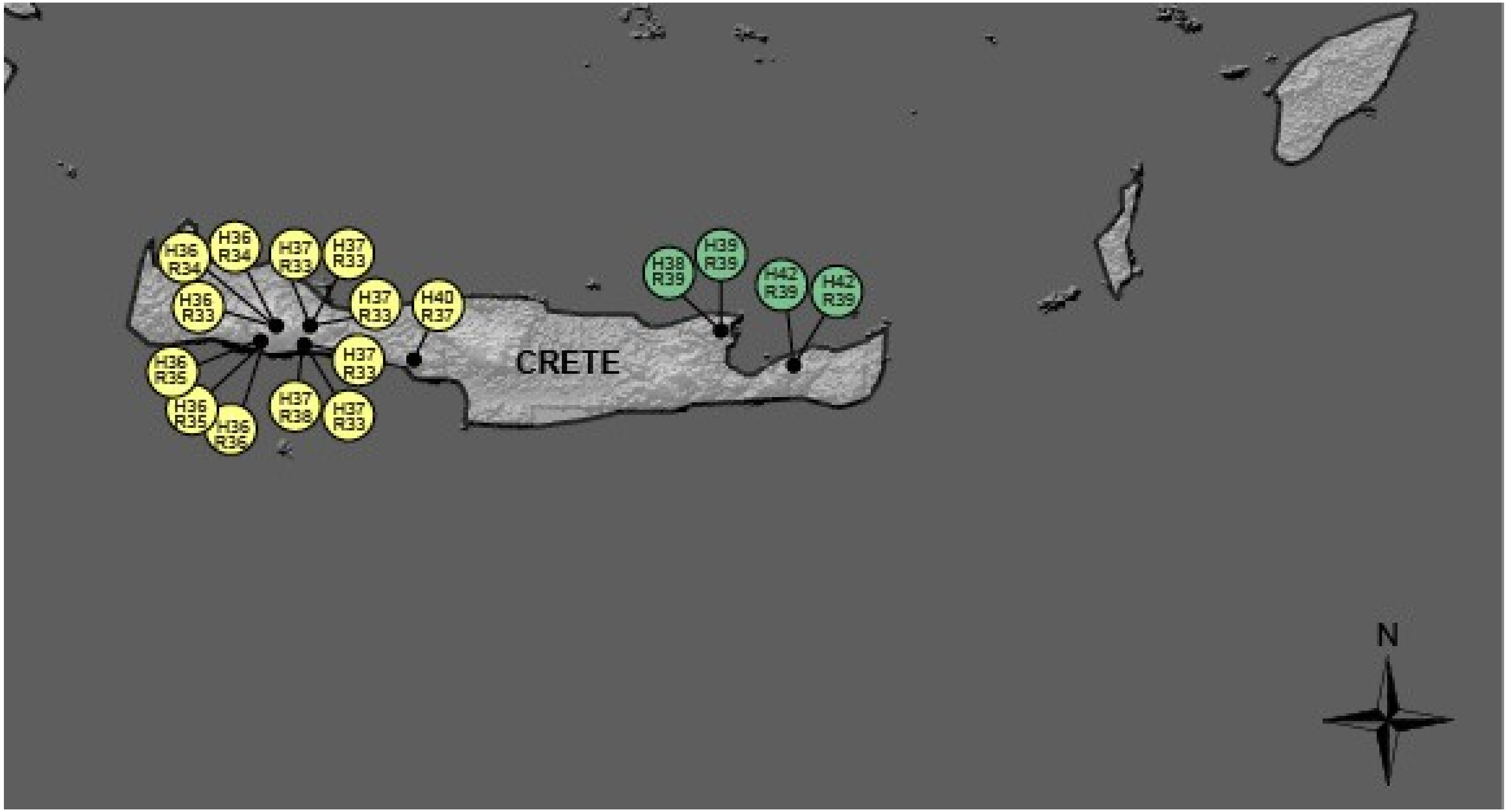


**Figure S3.** Spatial distribution of haplotypes and ribotypes in *Ricotia cretica*.

### Figure S4.pdf

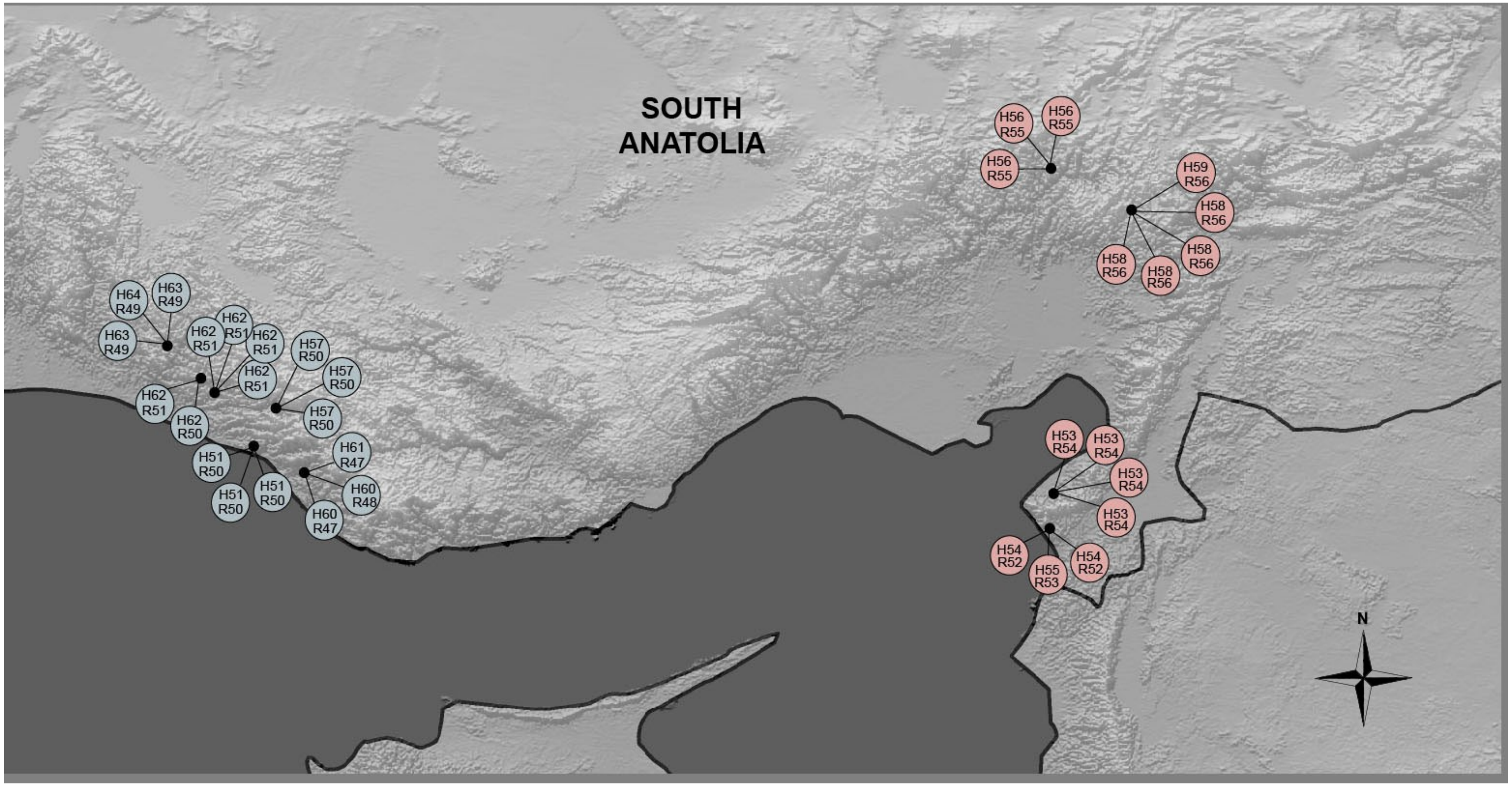


**Figure S2.** Spatial distribution of haplotypes and ribotypes in *Ricotia sinuata*.

### Figure S5.pdf

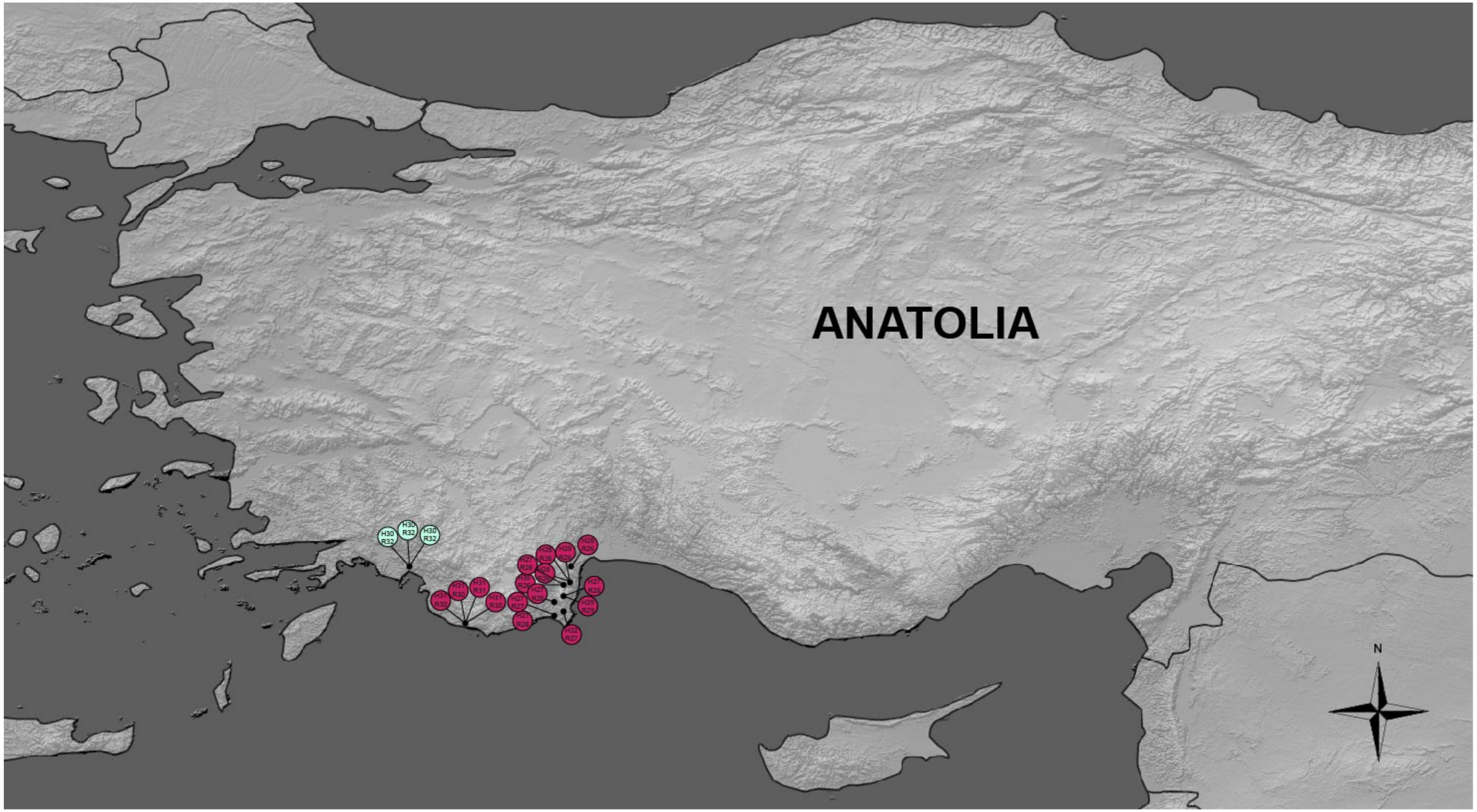


**Figure S2.** Spatial distribution of haplotypes and ribotypes in *Ricotia carnosula*.

### Figure S6.pdf

**Last Glacial Maximum (LGM) results of CCSM4 and MPI-ESM-P circulations models of *Ricotia* species**

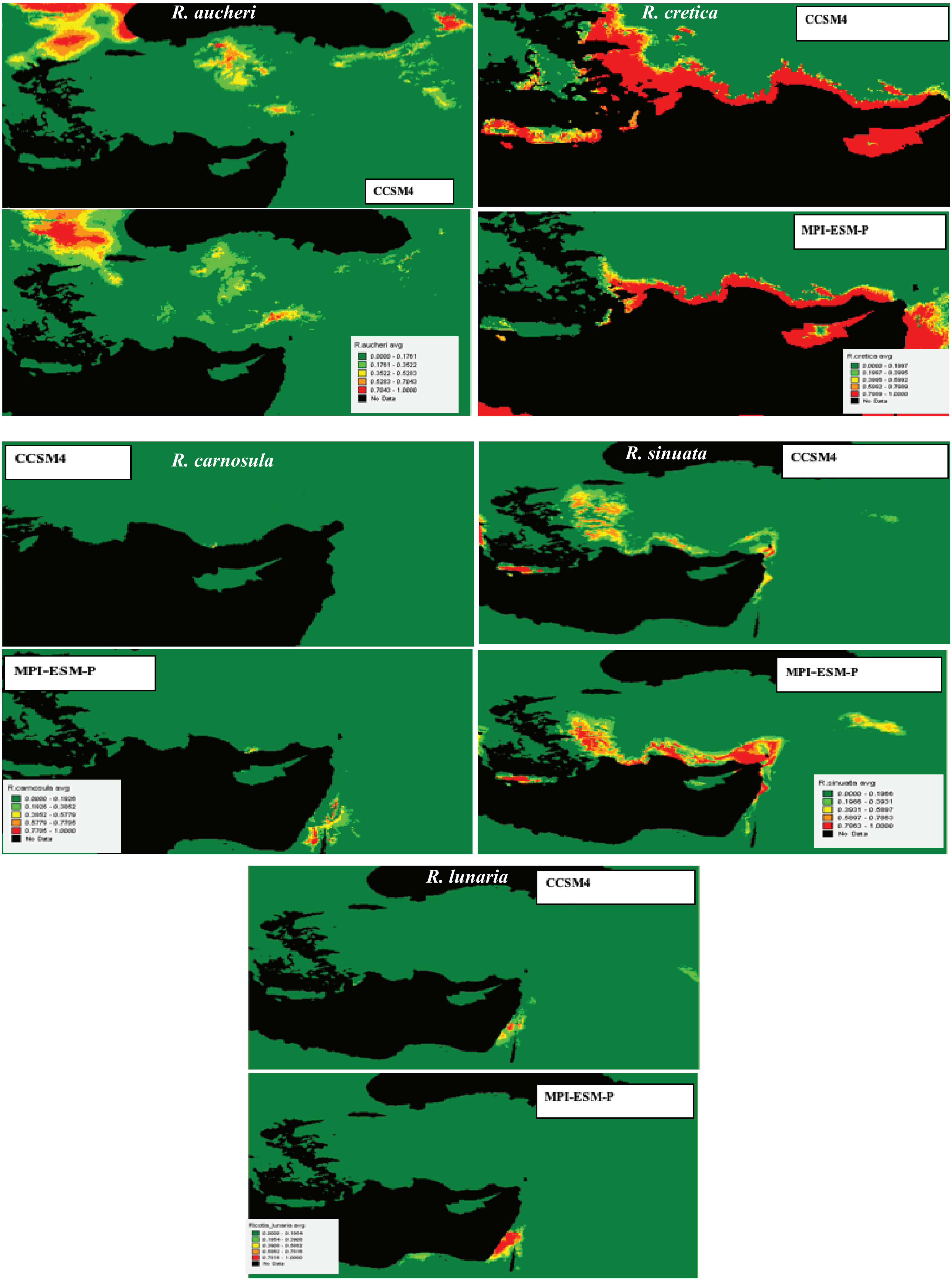
